# Functionally Distinct Subgroups of Oligodendrocyte Precursor Cells Integrate Neural Activity and Execute Myelin Formation

**DOI:** 10.1101/689505

**Authors:** Tobias Hoche, Roberta Marisca, Eneritz Agirre, Laura Jane Hoodless, Wenke Barkey, Franziska Auer, Gonçalo Castelo-Branco, Tim Czopka

**Affiliations:** Institute of Neuronal Cell Biology, Technical University of Munich, 80802 Munich, Germany; Munich Cluster of Systems Neurology (SyNergy), 81377 Munich, Germany; Graduate School of Systemic Neurosciences (GSN), Ludwig-Maximilian University of Munich, 82152 Planegg-Martinsried, Germany; Laboratory of Molecular Neurobiology, Department Medical Biochemistry and Biophysics, Biomedicum, Karolinska Institutet, Stockholm 17177, Sweden; Ming Wai Lau Centre for Reparative Medicine, Stockholm Node, Karolinska Institutet, 17177 Stockholm, Sweden

## Abstract

Recent reports revealed heterogeneity of oligodendrocyte precursor cells (OPCs). It remains unclear if heterogeneity reflects different types of cells with distinct functions, or rather transiently acquired states of cells with the same function. By integrating lineage formation of individual OPC clones, single-cell transcriptomics, calcium imaging and manipulation of neural activity, we show that OPCs in the zebrafish spinal cord can be divided into two functionally distinct entities. One subgroup forms elaborate networks of processes and exhibits a high degree of calcium signalling, but infrequently differentiates, despite contact to permissive axons. Instead, these OPCs divide in an activity and calcium dependent manner to produce another subgroup with higher process motility and less calcium signaling, which readily differentiates. Our data show that OPC subgroups are functionally diverse in responding to neurons and reveal that activity regulates proliferation of a subset of OPCs that is distinct from the cells that generate differentiated oligodendrocytes.

## Introduction

In the central nervous system (CNS) of vertebrates, oligodendrocyte precursor cells (OPCs, also referred to as NG2 cells) comprise an abundant cell population that tiles the CNS lifelong by forming an elaborate process network ^1^. OPCs are a major proliferative cell type in the CNS and are the cellular source for new myelin during CNS development, in response to neuronal activity in the context of myelin plasticity, and during regeneration of damaged myelin ^2-5^. We have a relatively robust understanding of the cell intrinsic signalling cascades and transcriptional changes that govern the differentiation of OPCs to myelinating oligodendrocytes ^6,7^. However, the CNS comprises many more OPCs than will ever differentiate. Therefore, a major question is if all OPCs can equally contribute to myelin formation, or if subsets of OPCs exist, with different fates and functions.

Various attempts to compartmentalize OPC properties have revealed that that these cells are indeed not a uniform population with equal properties ^8-13^. OPCs in different regions of the CNS show different responsiveness to growth factors ^14^, vary in their capacity to differentiate when transplanted into other CNS areas ^15^, and even disease-specific OPC phenotypes have been identified in mouse models of multiple sclerosis (MS) as well as human MS patients ^16,17^. Furthermore, it was recently shown that physiological properties of OPCs become increasingly diverse over time ^18^. Despite all these studies, however, a major remaining issue is that it is not clear whether the reported diversity of OPC properties represent bona fide types of OPCs with distinct functions, or rather different states of cells with the same function as they progress along their lineage, or while they reside in a particular microenvironment. The reason for this is that it is inherently difficult to definitively monitor the dynamics of cell fate, lineage progression and function from single time-point analyses, including those of sequencing datasets. So far, no study has carried out a systematic analysis of cell dynamics within the oligodendrocyte lineage over time, whilst probing the function and molecular states of subsets of OPCs in vivo. Such an analysis would help advance our understanding of how oligodendrocyte lineage cells relate to one another and if OPCs with different functions exist. It would, for example, help resolve open questions as to how neuronal activity affects OPC fates. OPCs integrate neuronal activity through their intimate contact with surrounding axons by virtue of the expression of a wide range of neurotransmitter receptors and voltage gated ion channels ^19-24^. However, the role of neural activity in controlling OPC fate decisions is still unclear, because increased neuronal activity increases both proliferation of OPCs and their differentiation into myelinating oligodendrocytes ^2,3,25-27^. This may be because subgroups of OPCs exhibit different electrophysiological properties and therefore differ in their sensitivity to axonal signals, or because they show differential cell fates in responses to increased activity. In any case, analysis of cell physiology alone, as well as analysis of gene expression or cell fate alone are not sufficient to reveal direct causality between heterogeneity in OPC properties and fates.

Here, we have addressed the diversity of OPC fates in an integrated approach by combining single-cell RNA sequencing in zebrafish, in vivo live cell imaging of properties and fates of distinct OPC clones, calcium imaging at single cell and population level, and manipulation of physiology to reveal how neural activity affects OPC subgroups in their ability to divide and to differentiate.

## Results

### OPCs form a network throughout the CNS that is composed of cells with distinct morphologies and dynamics

We have generated transgenic zebrafish lines that specifically label oligodendrocyte precursor cells (OPCs) using *olig1* upstream regulatory sequences (Fig. 1, Movie S1) ^28,29(and this study)^. Whole animal and high resolution imaging of Tg(olig1:memEYFP), Tg(olig1:nls-mApple) animals showed that labelled cells form a complex network of cellular processes that extends throughout the CNS, and which persists from late embryonic stages until at least the end of our analysis by three weeks of animal age (Fig. 1A, Movie S1). Cross-sectional views at the level of the spinal cord revealed that olig1:memEYFP labelled processes were almost exclusively found within axon-rich areas of the lateral spinal cord where myelinated and unmyelinated axons reside, and not in the neuron-dense regions of the DAPI-labelled medial spinal cord, as shown by co-labelling with the axon marker 3A10 and Tg(mbp:EGFP-CAAX) labelling of myelin (Fig. 1B, S1A). Quantitative analysis of cell numbers Tg(olig1:nls-mApple), Tg(mbp:nls-EGFP) animals to label OPCs and myelinating oligodendrocytes over the first month of animal development showed that the number of spinal cord OPCs remained stable over the entire period of analysis (mean of 29.6±6.4 cells/field at 3dpf *vs*. 32.0±5.9 cells/field at 28dpf, n=17/13 animals), while the number of myelinating oligodendrocytes steadily increased during the same time (mean of 7.3±5.7 cells/field at 3dpf *vs.* 95.2±13.4 cells/field at 28dpf, n=17/13 animals; Fig. 1C, S1B).

**Figure 1:**
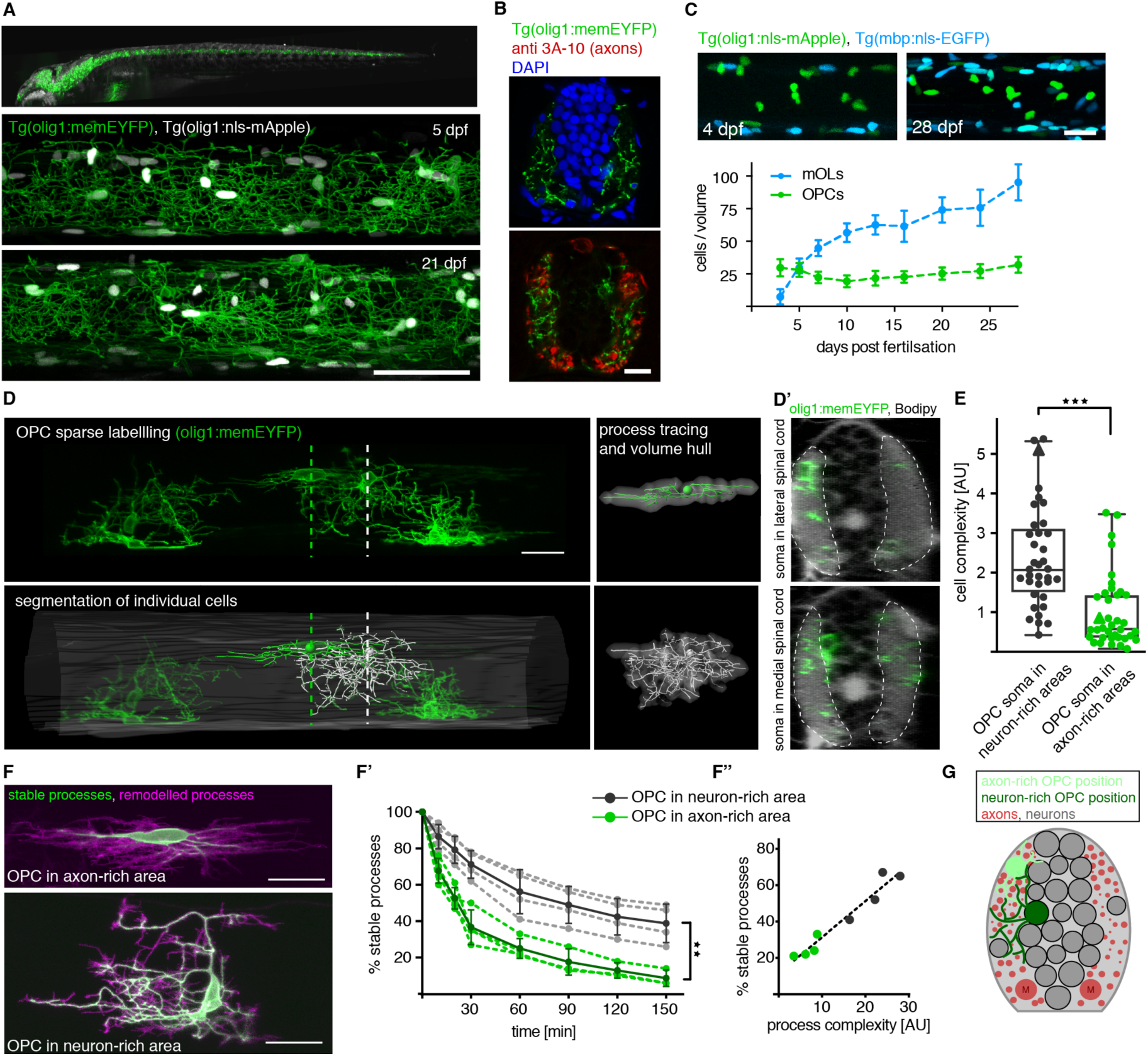
Characteristics of oligodendrocyte precursor cells (OPCs) in zebrafish. **A)** Top panel: Lightsheet microscopy image of whole Tg(olig1:memEYFP) transgenic animal at 5 days post-fertilization (dpf). Middle and bottom panels: confocal images of Tg(olig1:memEYFP), Tg(olig1:nls-mApple) zebrafish at the level of the spinal cord at 5 dpf and 21 dpf. Scale bar: 50 *µ*m. **B)** Cross-section of a Tg(olig1:memEYFP) spinal cord at 5 dpf stained for axons (3A-10 antibody) and nuclei (DAPI). Scale bar: 10 *µ*m. **C)** Top panel: Confocal images of Tg(mbp:nls-EGFP), Tg(olig1:nls-mApple) transgenic animals at 4 and 28 dpf. Below: Quantification of cell numbers of OPCs (olig1: nls-mApple-pos., mbp:nls-EGFP-neg.) and myelinating oligodendrocytes (mbp:nls-EGFP-pos.) in the spinal cord between 3 and 28 dpf. Data are expressed as mean ± SD, n(timepoint)=17/15/15/16/17/17/20/12/13. Scale bar: 20 *µ*m. **D)** Reconstruction of cell morphologies of individually labelled olig1:memEYFP cells. Three-dimensional rotations of olig1:memEYFP-labelled cells with a BODIPY counterstain to reveal the position of OPC somata within the tissue. Scale bar: 20 *µ*m. **E)** Quantification of relative cell complexities of olig1:memYFP labeled cells with their soma in neuron-rich and axon-rich areas of the spinal cord. Triangles indicate the cells shown in D. Data are expressed as median ± 25/75% interquartile percentiles (2.1±1.5/3.2 for OPCs in neuron-rich areas *vs*. 0.6 ±0.4/1.5 in axon-rich areas, n=36/38, p<0.001 (Mann-Whitney U test). **F)** Remodelling dynamics of individual olig1:memYFP labeled cells. Left: Projections of 60 minutes time-lapse imaging (5 minutes intervals) to show remodelled (magenta) and stable (green) processes of OPCs in axon-rich (top) and neuron-rich (bottom) spinal cord). Scale bar: 20 *µ*m. F’) Quantification of stable processes over time. Dashed lines connect data points of individual cells. Data points connected by continuous lines represent means ± SD within the groups. N=4/4 cells, p(position) <0.01 (two-way repeated-measures ANOVA of time-points 0-60 min). F”) Correlation between process network complexity and remodeling dynamics (process stability) of individual OPCs (at 60 min: Pearson’s r=0.974, 95% CI 0.859 to 0.995, R^2^=0.949, p<0.0001, n=8). The dashed line indicates linear regression curve (y=0.01988*x+11.48). **G)** Schematic overview depicting the positioning of OPCs in the in the zebrafish spinal cord. See also Fig. S1, Movies S1-S6

In order to investigate how individual cells contribute to this OPC process network over time, we carried out OPC sparse labelling and morphometric three-dimensional analysis of individual cells (Fig. 1D, D’; S1C, D; Movie S2). Our analysis revealed that OPC somata resided either within the neuron-rich areas of the medial spinal cord, or within the axon-rich areas of the lateral and ventral spinal cord (Fig. 1D”). Despite these different soma positions, the processes of each cell extended into axonal territories of the lateral spinal cord, as observed in the full transgenic lines (Fig. 1B).

Detailed four-dimensional analysis of single cell morphology and process dynamics revealed that OPCs display distinct properties with regard to process branching complexity and remodelling dynamics that correlate with their position in the spinal cord. We determined OPC complexity as a measure of the volume occupied by any given cell, and the summed process length and branch number within this volume (Fig. S1D). OPCs with their soma within neuron-rich areas of the medial spinal cord formed a much more elaborate process network compared to OPCs with their soma and processes entirely residing within lateral axonal territories (median 2.1±1.5/3.2 arbitrary units (AU) for OPCs in neuron-rich areas *vs*. 0.6 ±0.4/1.5 AU in axon-rich areas, n=36/38 cells, p<0.001, (Mann-Whitney U test); Fig. 1E). Furthermore, process dynamics of individual OPCs occurred at highly different rates, depending on their position within the spinal cord. Time-projections of a 60 minute imaging period showed that OPCs with cell bodies in the axon-rich tracts remodelled the vast majority of their process network, whereas OPCs that reside with somas in the neuron-rich areas only remodelled their process tips whilst retaining stable major branches during the same time (mean 25.0±5.5% stable processes for OPCs in axon-rich areas *vs*. 56.3±12.2% for cells in neuron-rich areas at 60 min, n=4/4 cells; two-way repeated-measures ANOVA of time-points 0-60 min, p(position)<0.01; Fig. 1F-G’, Movies S3-S6). Together, these data show that OPCs form two distinct subgroups with different branching complexity and remodelling dynamics, which can be discriminated by the position of their cell body within the spinal cord.

### Single-cell RNA sequencing reveals distinct molecular signatures of OPC subgroups

As we could distinguish two OPC subgroups with different properties relating to soma position, morphology and process dynamics, we carried out transcriptome analysis of single OPCs using SmartSeq2 ^30^ in order to determine the molecular signatures that underlie the observed differences in OPCs (Fig. 2, S2). Clustering analysis of 310 cells that we isolated by fluorescence activated cell sorting of Tg(olig1:memEYFP) animals led to the identification of five major cell clusters (#1 - #5) that were of genuine oligodendrocyte lineage identity, based on the co-expression of the key lineage transcription factors *sox10, nkx2.2a*, and *olig2* (Fig 2B, S2D). We confirmed oligodendrocyte lineage identity of cells labelled in our transgenic lines by *in-situ* hybridization and immunohistochemistry on tissue sections of Tg(olig1:nls-mApple), Tg(mbp:nls-EGFP) animals and found that almost all OPCs detected in the transgene (olig1 reporter-pos./ mbp reporter-neg.) also co-expressed Sox10 and *nkx2.2a* (62/62 Sox10-pos. OPCs, n=4 animals; 135/139 *nkx2.2a* pos. OPCs, n=12 animals; Fig. 2C, D, S2D). Two additional sequencing clusters (#6 and #7) were negative for oligodendrocyte lineage markers and thus not investigated further (Fig. S2C, D). Among the five oligodendrocyte lineage clusters #1 to #4 expressed bona-fide markers of OPCs, such as *ppp1r14bb* (Fig 2B, S2D). In contrast, cluster #5 expressed myelin genes like *plp1a* and *mbpa* and hence represents myelinating oligodendrocytes (Fig. 2B, S2D). Therefore, clusters #1 to #4 likely represent the OPCs investigated in our light microscopic analysis.

**Figure 2:**
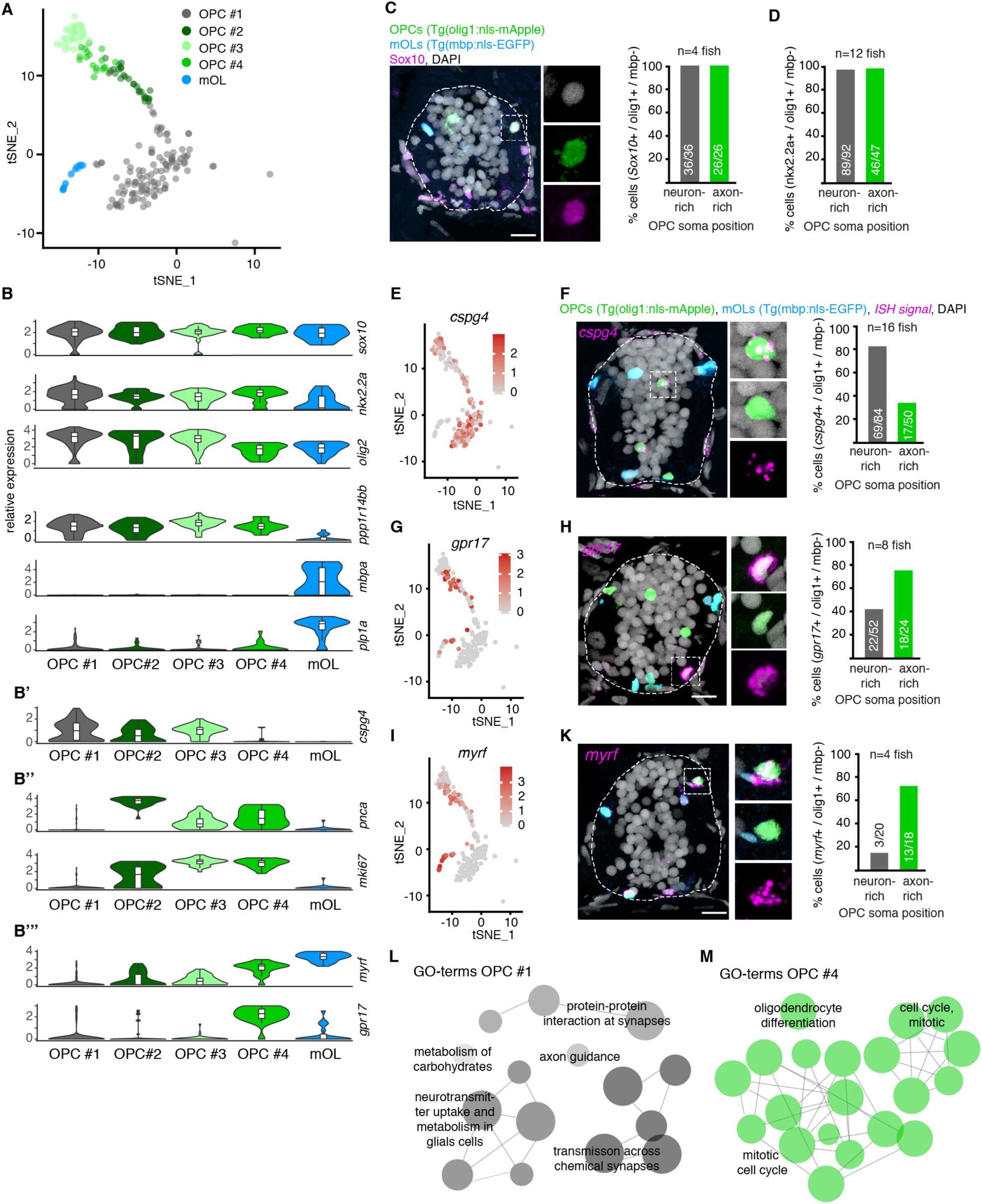
Single cell RNA sequencing of zebrafish OPCs. **A)** TSNE plots olig1:memYFP sorted cells with oligodendrocyte lineage identity. **B)** Violin plots with expression levels of key oligodendrocyte lineage markers in clusters and major myelin genes. Markers significantly enriched in clusters OPC #1 – 4 are shown under B’ – B”’. **C)** Immunohistochemical staining for Sox10 on transversal spinal cord sections of 7 dpf Tg(olig1:nls-mApple), Tg(mbp:nls-EGFP) animals and quantification of Sox10-expressing OPCs (olig1:nls-mApple-positive, mbp:nls-EGFP-negative) in neuron-rich and axon-rich areas, respectively. Scale bar: 10 *µ*m **D)** Quantification of *nkx2.2a-*expressing OPCs as described in C. Scale bar: 10 *µ*m. **E)** TSNE plots of *cspg4* expression levels in olig1:memEYFP-sorted cells. **F)** *In-situ* hybridization and quantification of *cspg4*-expressing OPCs as described in C. Scale bar: 10 *µ*m **G)** TSNE plots of *gpr17* expression levels in olig1:memEYFP-sorted cells. **H)** *In-situ* hybridization and quantification of *gpr17*-expressing OPCs as described in C. Scale bar:10 *µ*m **I)** TSNE plots of *myrf* expression levels in olig1:memEYFP-sorted cells. **K)** *In-situ* hybridization and quantification of *myrf*-expressing OPCs as described in C. Scale bar:10 *µ*m. **L)** Gene ontology (GO) terms of top 30 significantly expressed genes in cluster OPC #1. **M)** GO terms of top 30 significantly expressed genes in cluster OPC #4

In order to determine which of the four different OPC clusters identified by our transcriptomic analysis correspond to the OPC subgroups seen by live cell imaging, we carried out RNA *in-situ* hybridizations of candidate genes that showed strong differential expression between clusters and determined whether the respective cells were positioned with their soma in neuron-rich or axon-rich areas. Here, we found that the bona fide OPC marker *cspg4* was significantly enriched in cluster #1 (Fig 2B’, S2I), and predominantly expressed by OPCs (olig1:nls-mApple-positive, mbp:nls-EGFP-negative) positioned with their soma in neuron-rich areas of the medial spinal cord (82% (69/84 cells) in neuron-rich areas versus 34% (17/50 cells) in axon-rich areas, n=16 animals; Fig. 2E, F). Clusters #2 and #3 also expressed the OPC marker *cspg4*, but additionally expressed genes related to cell cycle progression. Here, cluster #2 contained genes specific for S-phase like *pcna*, whereas cluster #3 was enriched for genes specific for M-phase like *mki67* (Fig 2B”, S2E-H).

In contrast to clusters #1 to #3, cluster #4 expressed only low-levels of *cspg4*, but instead expressed genes associated with oligodendrocyte differentiation, such as *myrf* and *gpr17*, which could suggest a committed oligodendrocyte precursor (COP) identity (Fig 2B’, B”’, E, G, I) ^10,31,32^. However, in addition to these differentiation-related markers, cells in cluster #4 still expressed proliferation-related genes, meaning that these are not post-mitotic differentiating oligodendrocytes (Fig. 2B”, S2E, G). By *in-situ* hybridization for *myrf* and *gpr17* as genes enriched in cluster #4, we found that both of these markers predominantly labelled OPCs that reside within axon-rich areas of the lateral spinal cord (*gpr17*: 75% (18/24 cells) in axon-rich areas *vs.* 42% (22/52) in neuron-rich areas, n=8 animals; *myrf*: 72% (13/18) in axon-rich areas *vs.* 15% (3/20) in neuron-rich areas, n=4 animals; Fig. 2G-K). Thus, OPCs that reside with their soma in medial neuron-rich areas predominantly represent cells found in OPC cluster #1 (and potentially #2 and #3), whereas cells within axon-rich areas are enriched in cluster #4. Interestingly, gene ontology (GO) term analysis of genes enriched in cluster #1 comprised many terms involved in neurotransmitter sensing and calcium signaling, but lacked markers associated with oligodendrocyte differentiation (Fig. 2L, S2I). In contrast, OPCs in cluster #4 were enriched for genes involved in early differentiation, along with proliferation genes. Therefore, our data suggest that the subgroups of OPCs that reside with their soma in axon-rich and neuron-rich areas have different capacities to integrate signals from axons, which may lead to a differential ability to proliferate or to differentiate towards myelinating oligodendrocytes, respectively.

### Long-term fates and interrelationships between subgroups of OPCs

Having identified subgroups of OPCs with distinct behavioural characteristics and molecular profiles, we wondered if they represent functionally distinct entities. In order to investigate how the OPC subgroups relate to another and if they differentially contribute to myelination, we carried out clonal analyses of individual OPC fates. First, we carried out time-lapse imaging of OPCs that resided with their cell bodies either in the lateral axon-rich or medial neuron-rich areas between three and five days post fertilization (dpf). While we observed cell divisions by both OPC subgroups, the vast majority of OPCs (17/21) that resided within axon-rich areas started to differentiate and to ensheath axons within 24 hours of investigation, whereas none of the OPCs with their soma in the neuron-rich areas did so (Fig 3A, B, Movies S7-8). In order to test whether this finding was a transient / temporally restricted phenomenon, in which scenario the OPCs located in the neuron-rich areas would differentiate at later stages of animal development, we have carried out long-term analysis of 95 individual OPCs up until 14 dpf. A retrospective inspection of individual differentiated oligodendrocytes as determined by the formation of myelin sheaths showed that, independent of animal age, the vast majority of OPCs differentiated within axon-rich areas (95% (41/43) at 4dpf, 88% (36/41) at 7dpf, 82% (9/11) at 14dpf; Fig. 3C). Most of these much later differentiating cells also showed the characteristic morphology of fast remodelling OPCs with low process complexity, as we already observed during embryonic stages (Fig. S3A). Similarly, population analysis of all myelinating oligodendrocytes labelled in Tg(mbp:nls-EGFP) confirmed that the relative proportion of myelinating cells that reside with their soma in neuron-rich areas always remained a minority of no more than 11% of all oligodendrocytes throughout the time period of analysis (3.2±6.1% at 3dpf *vs.* 10.3±3.9% at 7dpf *vs.* 11.4±2.9% at 28dpf; n=17/15/13 animals; Fig. 3D).

**Figure 3:**
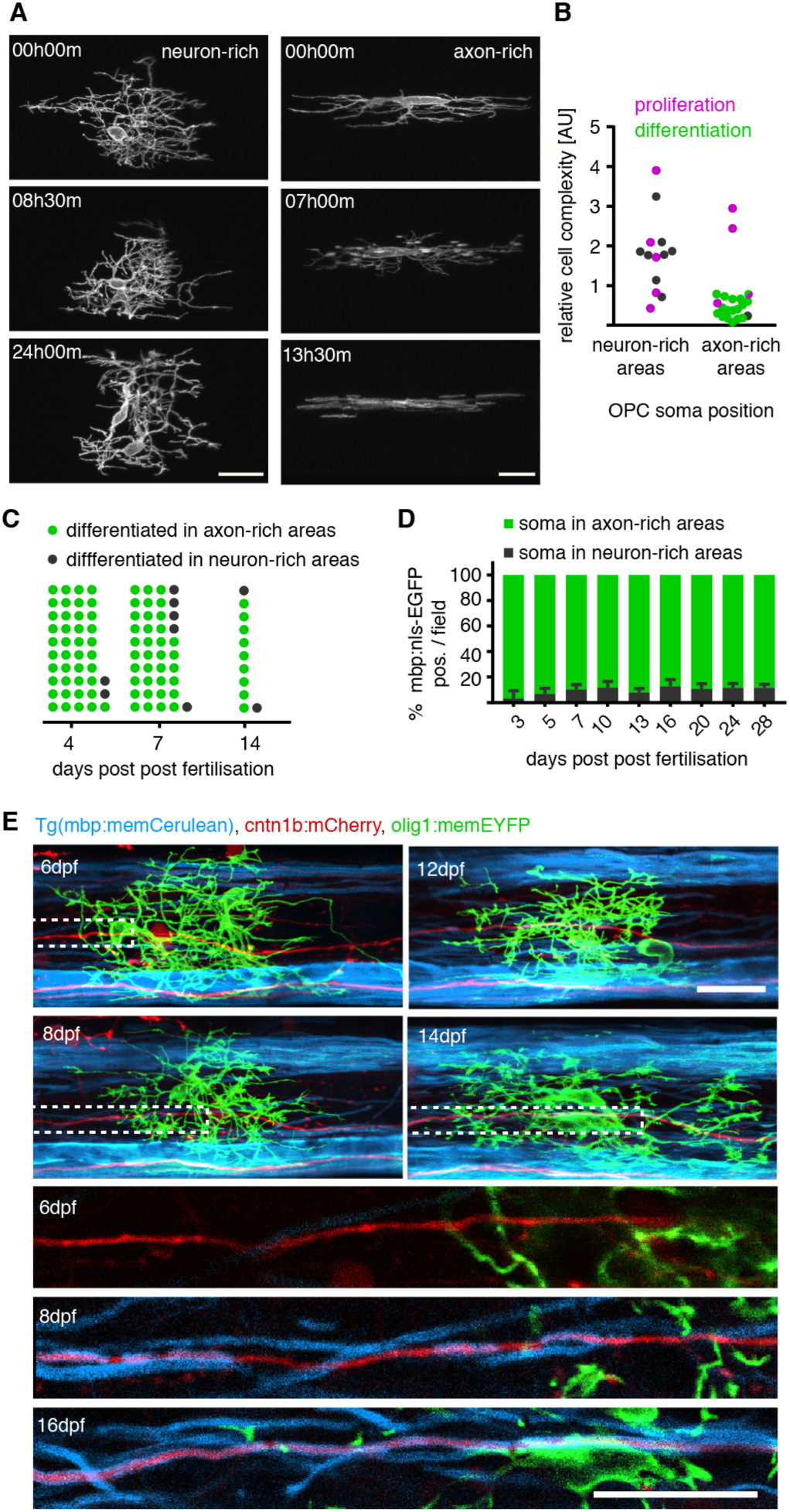
Differentiation properties of individual OPCs. **A)** Time-lapse imaging of individually labelled olig1:memEYFP cells with their soma in neuron-rich (left) and axon-rich (right) spinal cord at 3 dpf over up to 24 hours. Scale bar: 20 *µ*m. **B)** Quantification of proliferation / differentiation frequency and relative cell complexity of OPCs in neuron-rich and axon-rich areas over 24 h between 3 and 5 dpf. N=5/13 *vs.* 4/21 cells proliferated in neuron-rich *vs.* axon-rich areas, p=0.3 (Fischer’s exact test). N=0/13 *vs*. 17/21 cells differentiated in axon-rich vs neuron-rich areas, p<0.001 (Fischer’s exact test). **C)** Frequency distribution of OPCs positioning before differentiation at different developmental stages. **D)** Quantification of myelinating oligodendrocytes with their soma in neuron-rich areas in Tg(mbp:nls-EGFP) animals at different developmental stages. Data are expressed mean percentage ± SD. **E)** Time-lines of confocal images showing an olig1:memYFP labelled OPC with its soma in neuron rich areas between 6 and 16 dpf, while a cntn1b:mCherry labelled axon in close proximity becomes ensheathed with mbp:memCerulean labelled myelin. Scale bar: 20 *µ*m. See also Figure S3, Movies S7, S8

Here, it is important to note that (as described above) all OPC processes extend into the same axon-rich territories, regardless of the position of the respective cell body in neuron-rich or axon-rich areas (Fig. 1B, D”). Therefore, all OPCs can contact axons permissive for myelination, especially given that a single OPC can even span the entire dorso-ventral and medio-lateral dimensions of both spinal cord hemispheres (Fig. S1A, C, D). Indeed, using triple labelling of individual axons (cntn1b:mCherry) and OPCs (olig1:memEYFP) in a transgenic line that has all myelin labelled (Tg(mbp:memCerulean)), we were able to identify single OPCs that persisted undifferentiated from three until at least 16 dpf, while its processes were in close proximity to an axon that got increasingly myelinated during the same time (Fig. 3E). Therefore, these data show that OPCs that reside with their soma in neuron-rich areas can remain undifferentiated despite extending their processes into areas with target axons that are permissive for myelination.

The finding that one subset of OPCs (OPC#4) frequently differentiates while the other one rarely does (OPC#1), raised the question how these two groups relate to another. Where do new myelinating cells come from to prevent depletion of the respective pool of OPCs? One possibility would be that the cell body of slowly remodelling/ high complexity OPCs in the medial neuron-rich areas migrate into lateral axon-rich areas, where they would acquire a phenotype of fast remodelling myelinating OPCs typically found within these regions. Alternatively, new myelinating OPCs could arise from cell divisions where the daughter cell acquires a new phenotype. Indeed, we could observe cell divisions of OPCs in neuron-rich areas, followed by the migration of daughter cells into axon-rich areas and subsequent differentiation (Fig 4A, S4A). To investigate how to what extent this phenomenon accounts for the generation of new myelinating oligodendrocytes, we followed the fate of 113 individual OPCs with their soma in neuron-rich areas (Fig 4B, S4B). Over a time period of four days of analysis, 72% of the starting OPCs that we analysed (81/113 cells) divided, the remaining 28% (32/113 cells) did not. None of the non-dividing cells directly differentiated to myelinating oligodendrocytes. In contrast, we did observe the formation of new myelinating oligodendrocytes from 51% of the starting OPCs that also proliferated (42/81 cells). From 79% of initially dividing OPCs (33/42), all daughter cells differentiated in both neuron-rich and axon-rich areas. From the remaining 21% of initially dividing cells that gave rise to myelinating oligodendrocytes (9/42), some daughter cells differentiated while others remained as OPCs.

**Figure 4:**
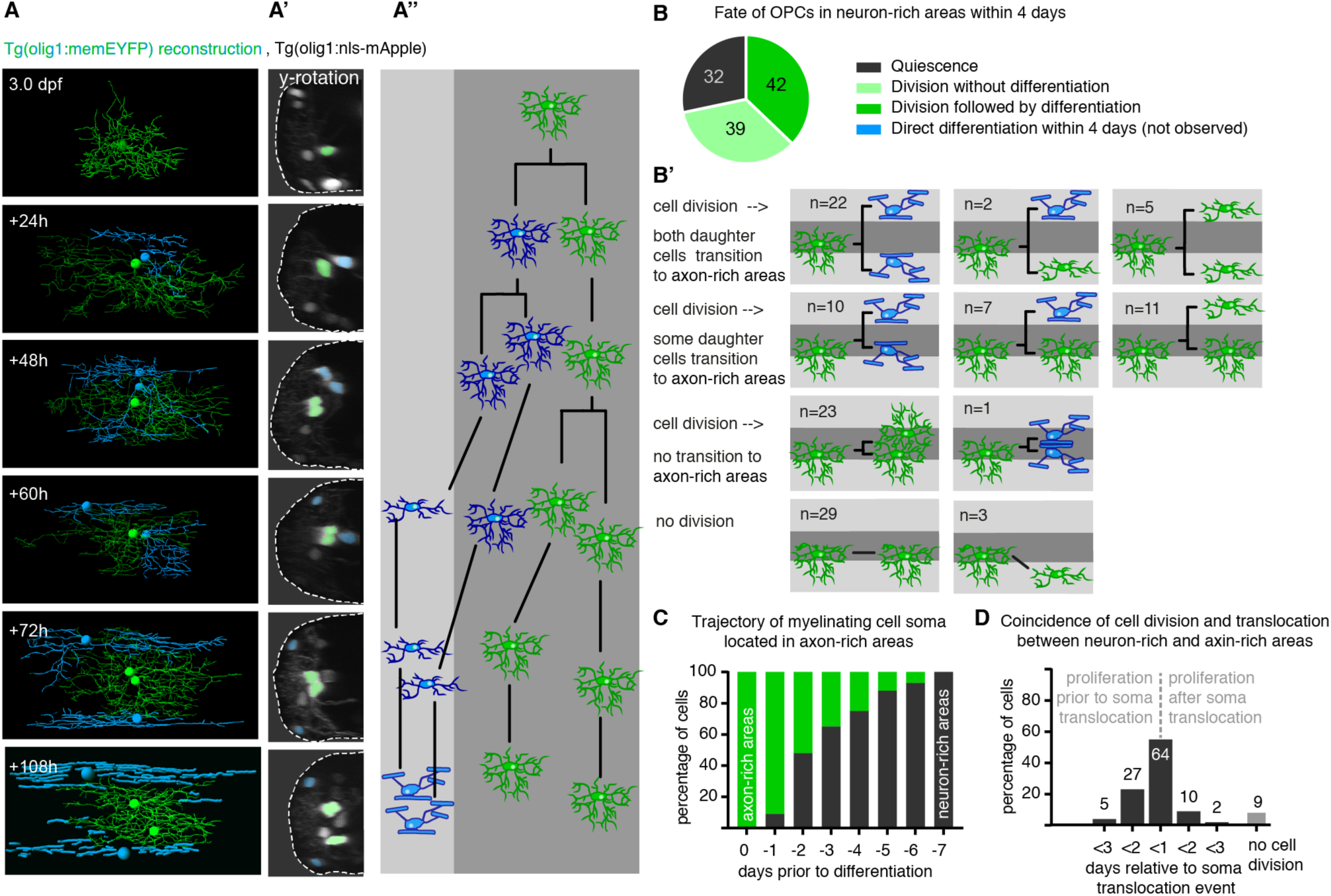
Differentiation properties of individual OPCs. **A)** Time-lapse imaging of individually labelled olig1:memEYFP cells with their soma in neuron-rich (left) and axon-rich (right) spinal cord at 3 dpf over up to 24 hours. Scale bar: 20 *µ*m. **B)** Quantification of proliferation / differentiation frequency and relative cell complexity of OPCs in neuron-rich and axon-rich areas over 24 h between 3 and 5 dpf. N=5/13 *vs.* 4/21 cells proliferated in neuron-rich *vs.* axon-rich areas, p=0.3 (Fischer’s exact test). N=0/13 *vs*. 17/21 cells differentiated in axon-rich vs neuron-rich areas, p<0.001 (Fischer’s exact test). **C)** Frequency distribution of OPCs positioning before differentiation at different developmental stages. **D)** Quantification of myelinating oligodendrocytes with their soma in neuron-rich areas in Tg(mbp:nls-EGFP) animals at different developmental stages. Data are expressed mean percentage ± SD. **E)** Time-lines of confocal images showing an olig1:memYFP labelled OPC with its soma in neuron rich areas between 6 and 16 dpf, while a cntn1b:mCherry labelled axon in close proximity becomes ensheathed with mbp:memCerulean labelled myelin. Scale bar: 20 *µ*m. See also Figure S3, Movies S7, S8

As a consequence, the majority of myelinating cells that are located in axon-rich areas originated from cell divisions of OPCs in neuron-rich areas (Fig 4C). These cell divisions occurred within two days around the soma translocation between neuron-rich and axon-rich areas (86% (101/117 cells); Fig 4C, D). From this cell fate analysis, we conclude that a hierarchy between the two OPC subgroups exists. OPCs with high process complexity and slow remodeling dynamics located in neuron-rich areas of the spinal cord will likely not differentiate directly, but they can divide to produce a daughter cell with higher cell motility and higher likelihood to differentiate to myelinating oligodendrocytes.

### Subgroups of OPCs show different degrees of calcium signaling activity

Because the population of infrequently differentiating OPCs (OPC #1) in the neuron-rich areas was strongly enriched in genes important for neurotransmitter signaling (Fig. 2L), we wondered if these cells communicate with axons in a distinct manner to the OPCs with cell bodies in the axon-rich areas that are more likely to myelinate. In such scenario, the rarely differentiating OPCs with high process complexity and rather slow remodelling dynamics might act as cohort of cells with functions to integrate signals from neurons. In order to test this, we generated new transgenic reagents and lines in which OPCs express the genetically encoded calcium sensor GCaMP6m, as it has previously been shown by several groups that OPCs respond to neural activity/neurotransmitter release with intracellular calcium rises ^21,33-35^ (Fig. 5). Fast (10Hz single plane) lightsheet imaging showed that GCaMP transients were detectable using olig1:GCaMP6m reporters which lasted between 2.2 and 3.6 seconds (average of 2.9s)(Fig. 5B). We assessed GCaMP signaling activity in individual OPCs expressing olig1:GCaMP6m-CAAX using three-dimensional time-lapse microscopy (Fig. 5A-E, Movies S9,S10). We could detect two different types of GCaMP transients. OPCs either showed transients that were restricted to process subdomains, which we observed in about 67% of cells (34/51) over a time period of 10 minutes of continuous imaging (Fig. 5C, E, Movie S9). In rare cases (3/51), however, GCaMP transients could spread throughout the entire cell (Fig. 5D, E, Movie S10). We also noticed that neighbouring OPCs did not necessarily show the same transients during the recording (Fig. 5C, D, Movies S9, S10), suggesting that individual OPCs may indeed show differential coupling to neurons. In order to investigate if subgroups of OPCs show differential calcium signalling activity, we generated a olig1:GCaMP6m full transgenic line to carry out population analysis of somatic calcium rises in volumes of several somites of whole spinal cord tissue (Fig. 5F, Movie S11). In our OPC calcium reporter animals, somatic OPC GCaMP rises could either remain restricted to single OPCs (Fig. 5F, G), affect groups of cells, and even all cells (Fig. S5). We then analysed GCaMP transients in all OPCs that were detectable in each sample. Interestingly, this revealed that in all animals investigated, OPCs within axon-rich areas had a lower probability to show GCaMP transients when compared to the OPCs that extend their processes into axon-rich areas, but which reside with their soma in the neuron-rich medial spinal cord (19% (9/48 cells) in axon-rich areas *vs*. 27% (73/285 cells) in neuron-rich areas, n=8 animals, p=0.007, paired Wilcoxon test; Fig 5G). Furthermore, the amplitudes of somatic GCaMP transients were also significantly lower in the subgroup of the OPCs within axon-rich areas (1.8±0.2 ΔF/F_0_ in axon-rich areas *vs*. 3.9±0.2 in neuron-rich areas, n=8 animals, p=<0.001, unpaired *t*-test; Fig. 5H).

**Figure 5:**
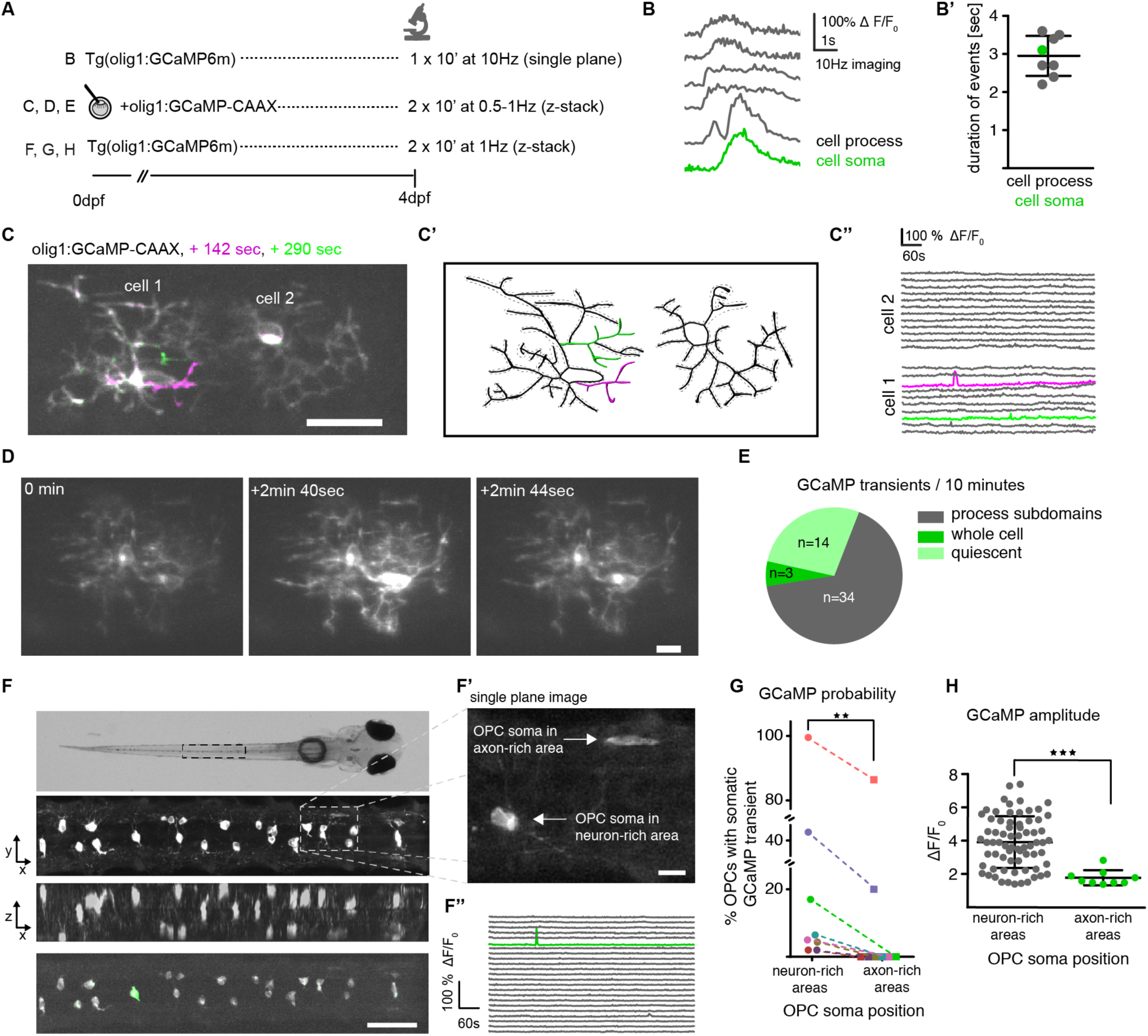
In vivo calcium imaging of individual OPCs. **A)** Overview of imaging conditions. **B)** Example traces of GCaMP transients in cell processes and soma of Tg(olig1:GCaMP6m) animals at four days post-fertilization (dpf). B’) Quantification of the duration of individual GCaMP transients. Data are expressed as mean ± SD. **C)** Projection of three timepoints (baseline: grey, t1: magenta, T2: green) of two individual olig1:GCamP-CAAX expressing cells showing transients restricted to process subdomains. Scale bar: 20 *µ*m. C’) Reconstruction of the cells shown in C. Active ROI are highlighted in green and magenta. C”) ΔF/F0 GCaMP transients of the different ROI shown in C’. **D)** Different time points of two individual olig1:GCaMP-CAAX expressing cells showing transients spreading throughout the cells. Scale bar: 10 *µ*m. **E)** Quantification of different types of GCaMP transients observed during an observation period of 10 minutes in individual olig1:GCamP-CAAX expressing cells. **F)** Top: Dorsal widefield view of a zebrafish at 4 dpf. Below: z-projection and z-rotation of Tg(olig1:GCaMP6m) in the spinal cord similar to as indicated by the boxed area above. Bottom: projection of 2 timepoints (baseline: grey, t1: green) showing a GCaMP transient restricted to a single cell in the volume. Scale bar: 50 *µ*m. F’) Single plane image taken from a z-stack of Tg(olig1:GCaMP6m) as indicated by boxed area showing individual OPCs with their soma in neuron-rich and axon-rich areas. Scale bar: 10 *µ*m. F”) ΔF/F_0_ GCaMP transients of individual cells shown in F (green trace depicts active soma). **G)** Quantification of probability of GCaMP transients in OPC somas in neuron-rich and axon-rich areas. N=7 animals, p=0.016 (Wilcoxon t-paired test). **H)** Quantification of GCaMP amplitudes measured in somata of OPCs in neuron-rich and axon-rich areas. N=8 animals, 81/10 cells, p=<0.001 (Welch’s t test). See also Fig. S5, Movies S9-S11.

In conclusion, the subgroup of OPCs that is less likely to differentiate directly, but which can divide to produce a myelinating daughter cell show higher rates of calcium signaling activity.

### Manipulation of neuronal activity and OPC calcium signalling mainly affects cell proliferation

If the subgroup of non-myelinating OPCs in the neuron-rich areas is most responsive to neuronal activity, as suggested by our GCaMP imaging data, we wondered how manipulation of neuronal activity affect Ca^2+^ activity, proliferation and differentiation behaviour of different OPC subgroups. We developed protocols that allowed us to enhance neural activity using bath application of the voltage-gated potassium channel blocker 4-Aminopyridin (4-AP) (Fig. 6A) – a drug that is also used to facilitate conduction in patients with Multiple Sclerosis ^36^. Single plane confocal imaging of OPC GCaMP transients in Tg(olig1:GCaMP6m) lines revealed that acute application of 0.5mM 4-AP enhanced the frequency of GCaMP transients in OPC processes within 15 minutes (median 12.5%±17.2 SD active ROI before 4-AP *vs*. 42.8%±25.4 SD after 4-AP, n=9 animals, p<0.01 (one-way ANOVA); Fig. 6B’). This increase was partially reversible by subsequent incubation with the voltage-gated sodium channel blocker Tetrodotoxin (TTX) (median 42.8%±25.4 SD 4-AP before TTX *vs.* 30%±25 SD after TTX, n=9 animals, p=0.36 (one-way ANOVA); Fig. 6B’). These manipulations suggest that a proportion of the increased OPC GCaMP transients induced by 4-AP are mediated by alterations in neuronal activity.

**Figure 6:**
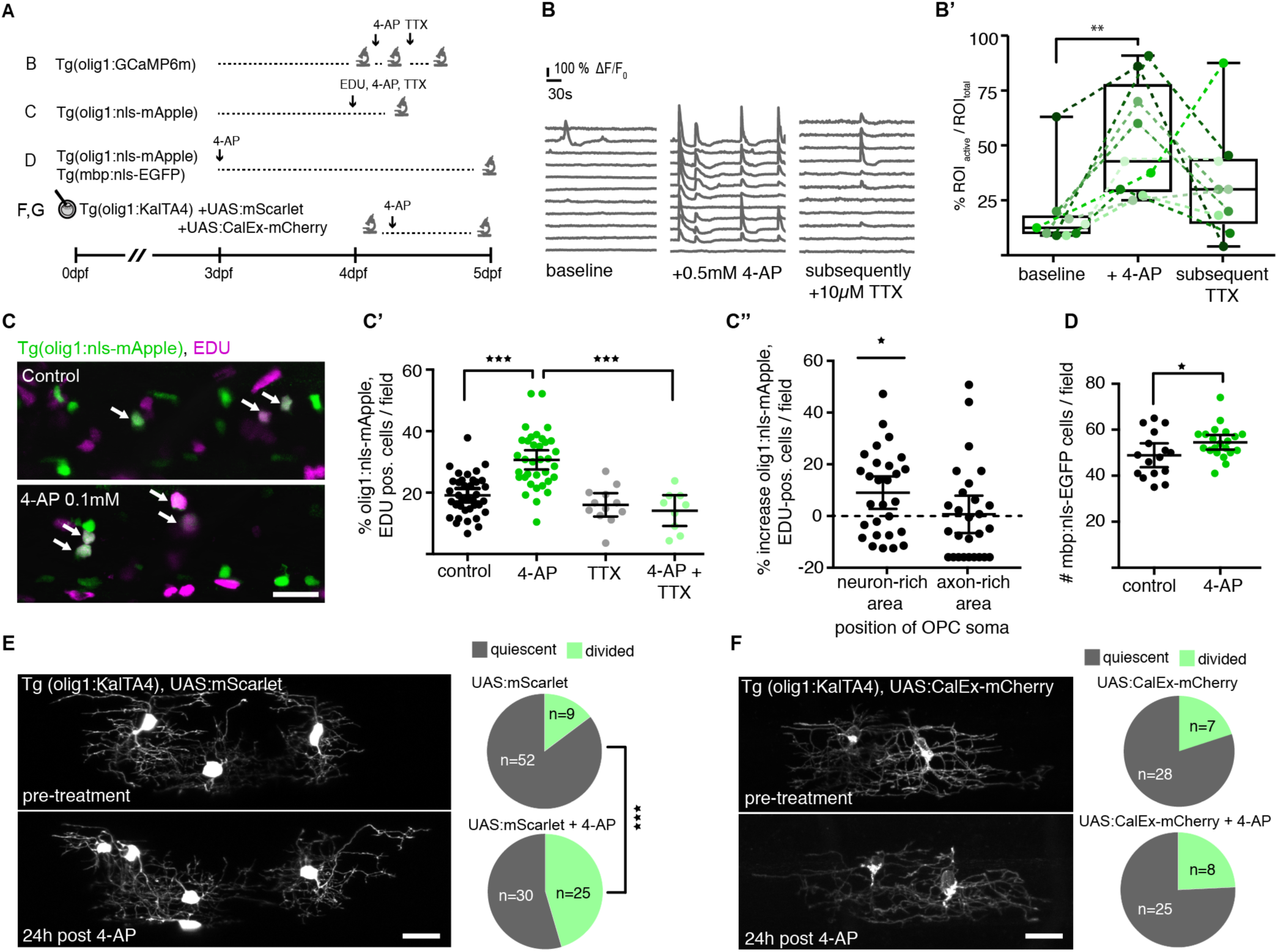
Manipulation of neural activity and calcium signalling in OPCs changes OPC proliferation. **A)** Overview of experimental paradigms. **B)** GCaMP traces in Tg(olig1:GCaMP6m) at 4dpf before (baseline) drug application, after treatment with 4-AP, and after addition of TTX to the same animal. B’) Quantification of GCaMP events (ROI active / ROI total) per fish in the different treatment conditions. Data are expressed at median ± 25/75% interquartile range, n=9 animals, p<0.01 (Kruskal-Wallis multiple comparison test). **C)** Confocal images of Tg(olig1:nls-mApple) zebrafish stained for EDU at 4dpf. Arrows indicate double positive cells. Scale bar: 50 *µ*m. C’) Quantification of olig1:nls-mApple/EDU double-positive cells in different treatment conditions. Data are expressed as mean ± 95% confidence interval, n=37/35/12/9 animals (control/4-AP/TTX/4-AP+TTX), p<0.001 (control *vs*. 4-AP and 4-AP *vs.* 4-AP+TTX), one-way ANOVA with Tukey’s multiple comparisons test. (C”) Quantification of increase in EDU-positive OPCs after 4-AP treatment located in neuron-rich and axon-rich areas. Data are expressed as mean ± 95% confidence interval, n=28/28 animals, p=0.02 (unpaired t-test). **D)** Quantification of mbp:nls-EGFP expressing cells with and without 4-AP treatment. Data are expressed as mean ± 95% confidence interval, n=16/21 animals, p=0.047 (unpaired t-test). **E)** Confocal images of Tg(olig1:KalTA4), UAS:mScarlet expressing OPCs at 4 dpf and 24h post 4-AP treatment. Scale bar: 20 *µ*m. Pie charts show the frequency of cell divisions observed in the different conditions. N=18/24 animals, p<0.001 (Fisher’s exact test). **F)** Confocal images of Tg(olig1:KalTA4), UAS:CalEx-mCherry expressing OPCs at 4 dpf and 24h post 4-AP treatment. Scale bar: 20 *µ*m. Pie charts show the frequency of cell divisions observed in the different conditions. N=11/24 animals.

In order to investigate how changes in neural activity and OPC calcium signalling affect cell behaviour, we developed a protocol for low-dose (0.1 mM), chronic 4-AP treatment for up to two days, which enhanced swimming behaviour (Fig. S6A) and GCaMP activity in neurons (Fig. S6B), without inducing detectable deleterious effects on tissue integrity, or causing inflammation in the spinal cord as assessed by macrophage recruitment (Fig. S6C).

In order to analyse how this manipulation affects OPC fate, we first assessed proliferation/S-phase entry by incorporation of EDU (Fig. 6C). Within 6 hours of 4-AP incubation, we detected a significant increase in EDU incorporation (mean 19.1±1.1% EDU^+^ OPCs in control *vs*. 30.6±1.5 in 4-AP, n=37/25 animals, p<0.001 (one-way ANOVA)), which was blocked in the presence of TTX (mean 30.6±1.5% EDU^+^ OPCs in 4-AP *vs.* 14.1±2.2% in 4-AP + TTX, n=12/9 animals, p<0.001 (one-way ANOVA); Fig. 6C’). This 4-AP induced increase in EDU-positive OPCs was mainly caused by cells that reside in neuron-rich areas (mean 19.4±2.0% EDU^+^ OPCs in control *vs*. 28.5±3.1% in 4-AP, n=28 animals, p=0.02 (unpaired *t*-test), whereas the number of EDU-positive OPCs within axon-rich areas remained unaltered (mean 15.9±2.8% EDU^+^ OPCs in control *vs*, 16.6±3.5% in 4-AP, n=28 animals, p=0.9 (unpaired *t*-test); Fig. 6C”). We could also detect a small increase in the number of myelinating oligodendrocytes in Tg(mbp:nls-EGFP) after two days of 4-AP incubation. This effect was, however, much less pronounced (mean 48.9±2.4 cells/field in control *vs*. 54.6±1.5 cells in 4-AP, n=16/21 animals, p=0.047 (unpaired *t*-test), Fig. 6D, S6D). In addition to EDU incorporation as a measure for S-phase entry, we also detected increased cell division rates of single UAS:mScarlet labelled OPCs in Tg(olig1:KalTA4) animals upon 4-AP incubation (15% (9/61 cells) in control *vs.* 45% (25/55 cells) in 4-AP, n=18/24 animals, p<0.001 (Fisher’s exact test); Fig. 6E).

Lastly, we tested whether intracellular calcium signaling is necessary for 4-AP mediated OPC proliferation. We expressed the calcium exporting pump hPMCA2w/b (referred to as CalEx) in order to reduce OPC calcium signaling, as was recently established in astrocytes ^37^. Single UAS:CalEx-mCherry expressing OPCs showed similar division rates as control UAS:mScarlet expressing OPCs (20% (7/35 cells) CalEx *vs*. 15% (9/61 cells) control, n=11/18 animals, p=0.6, Fisher’s exact test). However, in contrast to control OPCs, where 4-AP application triggered cell division (Fig 6E), UAS:CalEx-mCherry expressing OPCs did not show increased proliferation upon 4-AP incubation (23% (7/30 cells) control *vs*. 24% (8/33 cells) 4-AP, n=11/24 animals; Fig. 6F). Together, these date indicate that neural activity differentially affects the proliferation of distinct OPC subgroups, and that the OPC divisions triggered by 4-AP application require intracellular calcium signalling.

## Discussion

Oligodendrocyte precursor cells (OPCs) can divide or differentiate in response to appropriate stimuli to control oligodendrocyte cell numbers and to form new myelin. Using the spinal cord of larval zebrafish as model system, we have shown that OPC subgroups with distinct properties and functions exist. Our work provides novel insights into the heterogeneity of OPCs, the regulation of oligodendrocyte lineage formation, and the responses of OPC subgroups to neural activity.

The diversity of oligodendrocyte lineage cells is a topic of active research and various differences in oligodendrocyte properties have been assigned to subsets of cells, depending on regional origin, age, and local environment in the healthy and diseased brain ^8,38^. A major remaining question is, however, to what extent the observed differences between cells represent types of OPCs with different functions, or rather different states of cells with the same function, as they progress along their lineage or as they reside in a particular environment. Our results have revealed the existence OPC types with distinct functions. However, despite the different roles that the OPC populations identified here play within the same tissue to control division and differentiation, do they represent subgroups of intrinsically different cells? Our data do not argue for such genuine diversity, mostly because the determining criterium for the acquisition of the distinct functional properties seems to be the positioning of the cell soma. This is in line with previous studies which reported that OPCs are initially a rather homogenous population, despite their different developmental origins, and that diversification rather occurs as cells progress along their lineage ^10,12^. In the mammalian CNS, the most notable differences in OPC properties and differentiation capacities have been assigned to cells in the grey and white matter, which represent environments with differential permissiveness for differentiation, likely due to the different extent of available axons ^9^. In analogy to grey and white matter of the rodent CNS, the readily differentiating OPCs in the axon-rich areas of the zebrafish spinal cord that we investigated are probably very similar to white matter cells because these cells are entirely surrounded by axons. In contrast, however, the non-differentiating OPCs that reside with their soma in neuron-rich areas are not analogous to grey matter OPCs, because the cells in our system simultaneously extend their processes into axon-rich areas. This means that these OPCs are in the grey and white matter at the same time, but with different cellular compartments. This raises the question of which signals might induce the properties that the distinct OPC populations exhibit as they reside with their soma in neuron-rich and axon-rich areas, respectively. Furthermore, which signals trigger the differentiation of OPCs to myelinating oligodendrocytes if all OPCs contact the same cohort of axons? Axonal parameters such as their caliber, the presence/absence of inhibitory signals, or activity-dependent secreted factors are well-established in regulating axon ensheathment fate ^39^. However, our data argue that axonal signals are not sufficient to trigger OPC differentiation, a process that likely precedes axon choice for myelination by the same cell. Instead, OPCs with their soma in neuron-rich areas can divide to produce daughter cells with a much higher likelihood to differentiate to myelinating oligodendrocytes. The link between division and differentiation has previously been documented during mouse cortical development, where OPC differentiation is restricted to a short time window following its last division ^40,41^. Other studies, however, have reported direct differentiation of OPCs in the adult mouse cortex, particularly in the context of enhanced oligodendrogenesis in response to sensory enrichment and learning associated myelination ^2,4,27^. Whether these directly differentiating OPCs had divided just prior to analysis, or whether the directly differentiating OPCs in adulthood are slightly further along their lineage progression, remains to be determined. At the same time, although our data indicate that the likelihood to differentiate decreases as OPCs retain quiescent in the tissue, our study does not rule out that long-persisting OPCs can still differentiate.

In our work, we found that enhanced neural activity and OPC calcium signaling primarily lead to a rapid induction of cell division. The stimulatory effect of axonal activity on OPC proliferation has been observed in previous reports following optogenetic stimulation of axonal firing in mice, and in reverse by TTX injections to silence axons ^3,25^. However, we found that the OPCs that were most sensitive to neural activity were not the cells that likely differentiate to myelinating oligodendrocytes, but which instead divide in response to activity to create a daughter OPC with a much higher likelihood for differentiation. It is intriguing that these two OPC subpopulations also show highly different cellular properties: OPCs that effectively integrate neural activity form a much bigger process network and show slower process dynamics with major stable branches compared to the OPCs that readily differentiate, and which can remodel their entire process network within just a few minutes. Synaptic contacts have been described between axons and OPCs ^42^ and it would make sense for such synaptic contacts to be localised along OPC processes that remain stable for some time in order to form a structural synapse-like contact. In line with this thought, it has been reported that synaptic connections between axons and OPCs are rapidly lost as OPCs differentiate ^20,43^, and that axonal vesicle release regulates myelination via non-synaptic axon-OPCs contacts, at least *in vitro* ^44^. In the light of these published and our own findings, it is conceivable that synaptic axon-OPC contacts are not directly involved in regulating myelination (by e.g. serving as a signaling hub to define the site of future axon ensheathment), because the OPCs that form synapses with axons and might not be the same cells that will differentiate and ensheath axons. Why do OPCs integrate neural activity? From our data, it seems that one reason is to control their cell numbers. As a consequence, our data also indicate that a significant proportion of OPCs is not directly involved in cell differentiation to generate new myelin. This raises the question of how else OPCs might affect the brain, which remains to be addressed in future studies.

## Supporting information

Hoche Marisca et al supplementary figures

Hoche Marisca et al supplementary movie 1

Hoche Marisca et al supplementary movie 2

Hoche Marisca et al supplementary movie 3

Hoche Marisca et al supplementary movie 4

Hoche Marisca et al supplementary movie 5

Hoche Marisca et al supplementary movie 6

Hoche Marisca et al supplementary movie 7

Hoche Marisca et al supplementary movie 8

Hoche Marisca et al supplementary movie 9

Hoche Marisca et al supplementary movie 10

Hoche Marisca et al supplementary movie 11

## Acknowledgements

We are grateful to Rafael Almeida, David Lyons, Thomas Misgeld, Mikael Simons, and all members of the Czopka lab for their comments and suggestions during the assembly of the data and their presentation in this manuscript. We want to thank Baljit Khakh for the CalEx plasmid prior to publication, Kristen Kwan for providing pME_nls-Cerulean and pME_nls-mApple plasmids, and Douglas Kim for the GCaMP6m plasmid. We thank Single Cell Genomics Facility, Science for Life Laboratory, the National Genomics Infrastructure (NGI) and Uppmax for providing assistance in massive parallel sequencing and computational infrastructure. The bioinformatics computations were performed on resources provided by the Swedish National Infrastructure for Computing (SNIC) at UPPMAX, Uppsala University. E.A. is funded by European Union, Horizon 2020, Marie-Skłodowska Curie Actions, grant SOLO, number 794689. Work in G.C.-B.’s research group was supported by Swedish Research Council (grant 2015-03558), European Union (Horizon 2020 Research and Innovation Programme/ European Research Council Consolidator Grant EPIScOPE, grant agreement number 681893), Swedish Brain Foundation (FO2017-0075), Ming Wai Lau Centre for Reparative Medicine and Karolinska Institutet. Work in T.C.’s research group was funded by a Starting Grant of the European Research Council (ERC StG MecMy, grant agreement number 714440), the Deutsche Forschungsgemeinschaft (DFG, German Research Foundation) under its Emmy Noether programme for young investigators (ENP CZ226/1-1), and the DFG’s Excellence Strategy within the framework of the Munich Cluster for Systems Neurology (EXC 2145 SyNergy – ID 390857198).

## Author Contributions

Experiments were designed by T.H., R.M., L.J.H., W.B., G.C.B. and T.C.. Experiments were conducted by T.H., R.M., L.J.H., W.B. and F.A. Imaging data were analysed by T.H., R.M., L.J.H. and T.C.. E.A. and G.C.B. did the bioinformatic analysis of RNA sequencing data. T.C. conceived the project and wrote the manuscript with input from all authors.

## Methods Details

### Zebrafish lines and husbandry

We used the following existing zebrafish lines and strains: Tg(mbp:nls-EGFP) ^45^, Tg(mbp:EGFP-CAAX)^ue2Tg 46^, Tg(mbpa:MA-Cerulean)^tm101Tg 29^ (here referred to as Tg(mbp:memCerulean), Tg(elavl3:HSA.h2b-GCaMP6s)^jf5Tg 47^ (here referred to as Tg(elavl3:h2b-GCaMP6s), nacre and AB. The following lines were newly generated for this study: Tg(mfap:memCerulean), Tg(olig1:GCaMP6m), Tg(olig1:KalTA4), Tg(olig1:memYFP), Tg(olig1:nls-Cerulean) and Tg(olig1:nls-mApple). The olig1 gene regulatory sequence used drives reporter gene expression in OPCs; as OPCs differentiate to myelinating oligodendrocytes, reporter expression is downregulated. All animals were kept at 28.5 degrees with a 14/10 hour (h) light/dark cycle according to local animal welfare regulations. All experiments carried out with zebrafish at protected stages have been approved by the government of Upper Bavaria (animal protocols AZ55.2-1-54-2532-199-2015 and AZ55.2-1-54-2532-200-2015 to T.C.).

### Transgenesis constructs

Sequences for all primers used are listed in Table S1. In order to generate the middle entry clones pME_GCaMP6m, pME_GCaMP6m-CAAX and pME_mCherry-CalEx, the respective coding sequences were PCR amplified from template plasmids (pGP-CMV-GCaMP6m ^48^(gift from Douglas Kim & GENIE project, Addgene plasmid #40754); pZAC2.1_GfaABC_1_D mCherry-hPMCA2w/b ^37^ (gift from Baljit Khakh) and recombined with pDONR221 using BP clonase (Invitrogen). The middle entry clone pME_mScarlet was generated by BP recombination of mScarlet ^49^, of which the coding sequence with appropriate sites for recombination with pDONR221 was commercially synthesized by BioCat. The middle entry clone pME_KalTA4 has been published previously ^50^. The middle entry clones pME_nls-Cerulean and pME_nls-mApple were a kind gift of Kristen Kwan (Univeristy of Utah) ^51^. The 5’ entry clone p5E_mfap4 was generated by PCR amplification of a 1.5 kb DNA fragment of mfap4 upstream regulatory sequence ^52^ from AB genomic DNA and subsequent BP recombination into pDONRP4P1R (Invitrogen). The expression constructs pTol2_10xUAS:mScarlet, pTol2_10xUAS:CalEx-mCherry, pTol2_olig1(4.2):nls-Cerulean, pTol2_olig1(4.2):nls-mApple, pTol2_olig1(4.2):KalTA4 were generated using the entry clones described above and additional entry clones of the Tol2Kit ^51^ using Multisite LR recombination reactions. The expression constructs pTol2_olig1(4.2):memEYFP ^29^ and pTol2_cntn1b:mCherry ^53^ have been published previously.

### DNA microinjection for sparse labelling and generation of transgenic lines

Fertilised zebrafish eggs were microinjected with 1nl of an injection solution containing 5-25ng/µl DNA, 25-50ng/µl Tol2 transposase mRNA and 10% phenol red. Injected F0 animals were either used for single cell analysis, or raised to adulthood to generate full transgenic lines. For this, adult F0 animals were outcrossed with wildtype zebrafish and F1 offspring was screened for germline transmission of the fluorescent transgene.

### Pharmacological treatments

Zebrafish embryos at 3 and 4 days post fertilisation (dpf) were incubated in 4-aminopyridine (4-AP, Sigma-Aldrich) and/or Tetrodotoxin (TTX, Abcam) in Danieau’s solution. For long-term treatments (6-48 hours), 4-AP was used at 0.1mM and TTX was used at 50 *µ*m. For short-term treatments (<1 hour), 4-AP was used at 0.5mM and TTX at 10 *µ*m.

### Tissue preparation and cryosectioning for histology

5-7 days old zebrafish larvae were euthanised with 4mg/ml MS-222 and immersion fixed overnight at 4°C in 4% paraformaldehyde (PFA). Fixed animals were cryoprotected for a minimum of three days in an increasing concentration of sucrose (10%, 20%, 30%), embedded in TissueTek and stored at −80°C until sectioning. Transversal sections of 14-16 *µ*m thickness were cut using a Leica CM1850UV cryostat and subsequently stored at 80°C until further use.

### In-situ hybridization

RNA probes against zebrafish *nkx2.2a* (NM_001308640), *cspg4* (ENSDART00000112782), *myrf* (ENSDART00000157117), and *gpr17* (ACD product #504601) were purchased from ACD. We used the RNAscope Multiplex Fluorescent v2 kit (ACD) on cryosections according to the manufacturer’s protocol for fixed-frozen samples. Signals were detected using TSA-conjugated Opal dyes as listed in Table 2 (Perkin Elmer). After RNA hybridization, immunohistochemistry was carried out to detect transgenically expressed fluorescent proteins as described.

### Immunohistochemistry

All antibodies used are listed in Table S2. First, sections were blocked for 1.5 h at room temperature in PBS, 0.1% Tween20, 10% FCS, 0.1% BSA and 3% normal goat serum. Primary antibodies were incubated overnight at 4°C in blocking solution. Afterwards, sections were washed three times in PBS, 0.1% Tween20 and then incubated with appropriated Alexa Fluor-conjugated secondary antibodies (Invitrogen). Stained sections were washed two times in PBS, 0.1% Tween20 and once in PBS and subsequently mounted with ProLong Diamond Antifade Mountant with DAPI (Thermo Fisher Scientific). Sections were stored at 4°C.

### EDU incorporation assay

At 4 dpf Tg(olig1:nls-mApple) zebrafish embryos were incubated in 0.4 mM 5-ethynyl-2’-deoxyuridine (EDU) in Danieau’s solution. After 6 h incubation, embryos were incubated for 15 minutes in 2mg/ml Pronase (Sigma Aldrich) and subsequently fixed for 2h in 4% PFA. Whole embryos were stained for EDU using the Click-IT^™^ Alexa Fluor^™^ 647 Imaging Kit (ThermoFisher Scientific) as detailed in the kit protocol, with the exception of a 1.5h ClickIT reaction incubation time. Afterwards, immunofluorescence staining was performed for transgene detection.

### Mounting of embryonic and larval zebrafish for live cell microscopy

Animals were either anaesthetized with 0.2 mg/ml MS-222 (PHARMAQ, UK), or (in case of GCaMP imaging) immobilized with 0.5 mg/ml of the non-depolarizing neuromuscular junction blocker mivacurium chloride (Abcam). For confocal microscopy, animals were mounted laterally in 1% ultrapure low melting point agarose (Invitrogen) on a glass coverslip. The coverslip was flipped over on a glass slide with a ring of high-vacuum grease filled with a drop of Danieau’s solution to prevent drying out of the agarose. For lightsheet microscopy, embryos were mounted upright in low melting point agarose in a U-shaped glass capillary (Leica). After imaging, the animals were either sacrificed or released from the agarose using microsurgery blades and kept individually until further use.

### Confocal microscopy

12-bit confocal images were acquired on Leica TCS SP8 laser scanning microscopes. We used 405 nm wavelength for excitation of DAPI; 448 and 458 nm for excitation of Cerulean; 488 nm for EGFP and Alexa Fluor(AF) 488; 514 nm for EYFP; 552 and 561 nm for mApple, mScarlet, mCherry and AF555; 633 nm for AF633, AF647, Opal650 and BODIPY630/650. For fast confocal live cell imaging of cell motility and GCaMP transients, we used an 8 kHz resonant scanner. All other acquisitions were carried out with a Galvo scanner. For overview images and analysis of cell numbers (i.e. nuclear transgenes and EDU), we used 10x/0.4NA (acquisition with 568nm pixel size (xy), 2 *µ*m z-spacing) and 20x/0.7NA (acquisition with 142nm pixel size (xy), 1 *µ*m z-spacing) objectives. For analysis of stained cryosections, we used 63x/1.2NA H_2_O, and 63x/1.3NA glycerol objectives and acquired images with at least 100nm pixel size (xy) and 1 *µ*m z-spacing. For all other analysis, images were acquired using a 25x/0.95NA H_2_O objective with 114-151nm pixel size (xy) and 1 *µ*m z-spacing. When images were acquired for subsequent deconvolution, x/y/z parameters were increased closer to Nyquist resolution to be compatible for processing with Huygens software.

### Lightsheet microscopy

Lightsheet images were acquired at a Leica TCS SP8 DLS using a 2.5x/0.07NA illumination objective and 10X/0.3NA and 25X/0.95NA detection objectives with 2.5 mm deflection mirrors. Timelapses of GCaMP fluorescence were acquired using 488nm excitation wavelength. A ROI was drawn around either individual cells or a portion (5-7 somites) of whole zebrafish spinal cord tissue. Time-lapses of z-stacks were taken with a frame rate of 0.5-1Hz for 2×10 minutes, with a break of 10 minutes in between.

### Assessment of zebrafish swimming behaviour

Single 4dpf zebrafish were placed in a 3cm dish in 3ml Danieau’s solution (± 0.1mM 4-AP and/or 50 *µ*m TTX) and imaged for two minutes at 16 frames per second using a Hamamatsu Orca-05G equipped with a Kowa LM35JC10M objective.

### Fluorescence activated cell sorting of single zebrafish OPCs

Approximately 1000 Tg(olig1:memEYFP) at 5dpf were euthanised and de-yolked by repetitive pipetting embryos in de-yolking buffer (55mM NaCl, 1.2mM KCl, 1.25mM NaHCO3) with a P1000 pipette tip. Following two wash steps in Danieau’s buffer and centrifugation for 1’ at 300 g, tissues were digested for 30 minutes at 37°C in a shaking incubator using the Papain Dissociation Kit (Worthington Biochemical Corporations) according to manufacturer’s instructions. The obtained cell pellet was resuspended in FACSmax Cell Dissociation Buffer (Amsbio) with 5-10% FCS and filtered through a 30 *µ*m Filcon syringe (BD Bioscience) before sorting. Sorting of olig1:memEYFP cells followed a two-step protocol using a MoFlo XDP cell sorter (Beckman Coulter). First, approximately 100,000 EYFP-positive / propidium iodide (Thermo Fisher Scientific)-negative cells were sorted to exclude dead cells. The obtained cell suspension was then sorted again for EYFP-positive / Vybrant DyeCycle^™^ Ruby (Thermo Fisher Scientific)-positive cells. Single cells were sorted into each well of a 384-well plate containing RNA lysis buffer for Smart-seq2 (provided by ESCG of the Karolinska Institutet, Stockholm), as described in previously published work ^30^. Plates were stored at −80°C until further processing.

### Single-cell RNA sequencing

Single cell sequencing was performed on an Illumina HiSeq2500 instrument with the following specifications. The run was on a high-output flow cell with 50 bp single-read, clustered on a cBot with V4 kit. Reads were trimmed with Cutadapt 1.8.0 ^54^ and aligned to the reference genome GRCz11 using STAR 2.5.1.b ^55^ and using ENSEMBL94 transcript annotations, with the following parameters: --sjdbFileChrStartEnd SJ.out.tab --sjdbScore 0, --outFilterMatchNmin 10, --outSAMunmapped Within, -- quantMode TranscriptomeSAM, including the splicing junction database calculated by STAR on the same single cells in a first alignment. Aligned single cell reads were sorted and transformed to bam files using Samtools 1.3. Gene expression was calculated with Salmon 0.9.1 ^56^ using the sorted bam files as input; from the outputs we used the TPM gene expression values to build the expression matrix for the 384 cells. Cells were clustered with Seurat 3 ^57^ and filtered based on the distribution of gene expression (minimum 500 genes expressed per cell) and mitochondrial gene expression (maximum 0.05%). The remaining 310 cells were log-normalized individually with a scale of factor of 10,000. For downstream analyses we used the top 2000 variable genes. The shared-nearest neighbour (SNN) graph was constructed on cell-to-cell distance matrix from top 50 PCs. The SNN graph with resolution 1 was used as an input for the smart local moving (SLM) algorithm to obtain cell clusters, and visualized with *t*-distributed stochastic neighbour embedding (*t*-SNE). We identified 5 cell clusters where significantly differentially expressed genes (Wilcoxon rank sum test, min.pct = 0.25, thresh.use = 0.25, test.use = “wilcox”) list for each of the clusters allowed us to identify 2 VLMC clusters, 1 OL cluster, 1 OPC cluster and 1 COPs cluster. Specific markers for visualization on the t-SNE allowed us to identify 3 subclusters of COPs, that were later subset based on gene marker expression.

We retrieved all cell cycle genes from Gene Ontology annotations from Biomart and the human gene lists from Seurat v3 of S phase and G2m phase genes. First, the human annotations were transformed to ortholog gene name symbols to zebrafish using Biomart api. Then, we identified the PC containing cell cycle genes, this resulted mainly in PC2 and PC4. The cell cycle genes signal was also present in other PCs and for this reason we decided to regress out all cell cycle gene expression from the raw expression matrix. Comparison between cell cycle regress and non-regress did not lead on change on the main cell type clusters.

GO Term analysis of the different clusters was performed with ClueGo ^58^ using the top 30 upregulated genes within each cluster compared to all other clusters. We selected the following parameters: significant terms p<0.05 (Right-sided hypergeometric test for Enrichment, Benjamin-Hochberg p value correction) from the following databases: GO/Biological Processes (27-02-2019), GO/Molecular_Function (27-02-2019), Reactome/Pathways (27-02-2019) for all GO Tree intervals using GO term fusion. Genes from each cluster must represent at least 1% of all genes within a GO term (with a minimal absolute number of n=2 for cluster OPC #1, n=3 for OPC #4, n=6 for OPC #2, and n=7 for OPC #3) to be assigned to this cluster. The kappa score was always set to 0.4.

### Analysis and presentation of imaging data

All data were analysed using Fiji ^59^, Imaris 8.4.2 (Bitplane), Huygens Essential, Matlab, Microsoft Excel, Graphpad Prism, Adobe Photoshop and Illustrator. The Imaris Filament Tracer was used for three-dimensional reconstructions of OPC morphology. First, by tracing of the entire process network of individual OPCs we obtained information on total process length and branch point number of a cell. Second, we constructed a 3-dimentional hull around the filament network to obtain a measure for the volume of a cell using the Surface Tool in Imaris. Cell complexity was expressed as product of branch point density (branch point number divided by summed process length) and volume.

To analyse process remodelling dynamics of individual cells, cumulative maximum intensity time-projections of inverted greyscale images of OPC timelapses were generated to obtain a measure of the percentage of stable cell proportion at each time point relative to the starting time point. Measurements were done using the FIJI-plugin NeuronJ ^60^.

To trace GCaMP transients, regions of interest (ROI) were drawn using the Fiji ROI manager around the somata and/or processes of single cells. Traces of individual ROI are shown as ΔF/F_0_ by expressing the maximum fluorescent intensity at each time point normalized to the first 100 frames of each ROI using a custom written Matlab script. GCaMP transients were counted as events when they went beyond 40% ΔF/F_0_ for the frequency of somatic transients, and 30% ΔF/F_0_ for all other analysis. The duration of GCaMP transients is given as the half-width of the maximum ΔF/F_0_ for each event. To analyse changes in GCaMP frequency after pharmacological treatments, we used mean intensity of GCaMP fluorescence traces in ROI that were drawn around the same cells/processes in each treatment condition (baseline, after 4-AP, after TTX). For each ROI, fluorescence was normalized to the first 100 frames, and GCaMP events were determined using a threshold of 30% above average fluorescence change using the spike detrend function (https://de.mathworks.com/help/finance/tsmovavg.html).

Statistical analysis was done with Microsoft Excel and Graphpad Prism. Data were prepared for the figures with Fiji, Imaris 8.4.2, Graphpad Prism, Adobe Photoshop and Illustrator. All data were tested for normal distribution using the Shapiro-Wilk normality test. In the figures, normally distributed data are shown as mean ± standard deviation (SD) or 95% confidence intervals (CI), whereas non-normally distributed data are given as median with the 25% and 75% percentiles. For better readability of the main text, mean and median values are sometimes given with SD or CI as indicated, independent of normality. For statistical tests of normally distributed data comparing two groups we used unpaired and paired t tests. Non-normally distributed data were tested for statistical significance using Mann-Whitney U test (unpaired data) and Wilcoxon signed-rank test (paired data). To compare multiple (>2) groups ANOVA was used in combination with Tukey’s (paired data) or Kruskal-Wallis (unpaired data) multiple comparisons test. To analyse contingency tables, we used Fisher’s exact test. Statistical significance is given as p-value in the main text and figure legends, and indicated as *(p < 0.05), **(p < 0.01), ***(p < 0.001) in the figures.

## Data availability

Raw sequence data, gene expression and cell type annotation tables have been deposited in the Gene Expression Omnibus (GEO) under accession number <u>GSE132166</u>. A browsable webresource is available at https://castelobranco.shinyapps.io/zebrafish_OPCs/

## References

1. Oligodendrocyte Development and Plasticity. Cold Spring Harb Perspect Biol 8, a020453 (2015).

2. Hughes, E. G., Orthmann-Murphy, J. L., Langseth, A. J. & Bergles, D. E. Myelin remodeling through experience-dependent oligodendrogenesis in the adult somatosensory cortex. Nat Neurosci 21, 696–706 (2018).

3. Gibson, E. M. et al. Neuronal Activity Promotes Oligodendrogenesis and Adaptive Myelination in the Mammalian Brain. Science 344, 1252304 (2014).

4. McKenzie, I. A. et al. Motor skill learning requires active central myelination. Science 346, 318–322 (2014).

5. Zawadzka, M. et al. CNS-resident glial progenitor/stem cells produce Schwann cells as well as oligodendrocytes during repair of CNS demyelination. Cell Stem Cell 6, 578–590 (2010).

6. Emery, B. & Lu, Q. R. Transcriptional and Epigenetic Regulation of Oligodendrocyte Development and Myelination in the Central Nervous System. Cold Spring Harb Perspect Biol 7, a020461 (2015).

7. Liu, J., Moyon, S., Hernandez, M. & Casaccia, P. Epigenetic control of oligodendrocyte development: adding new players to old keepers. Current Opinion in Neurobiology 39, 133–138 (2016).

8. Foerster, S., Hill, M. F. E. & Franklin, R. J. M. Diversity in the oligodendrocyte lineage: Plasticity or heterogeneity? Glia 25, 2411 (2019).

9. Viganò, F. & Dimou, L. The heterogenic nature of NG2-glia. Brain Res. 1368, 129–137 (2015).

10. Marques, S. et al. Oligodendrocyte heterogeneity in the mouse juvenile and adult central nervous system. Science 352, 1326–1329 (2016).

11. Emery, B. & Barres, B. A. Unlocking CNS cell type heterogeneity. Cell 135, 596–598 (2008).

12. Marques, S. et al. Transcriptional Convergence of Oligodendrocyte Lineage Progenitors during Development. Developmental Cell 46, 504–517.e7 (2018).

13. Richardson, W. D., Young, K. M., Tripathi, R. B. & McKenzie, I. NG2-glia as Multipotent Neural Stem Cells: Fact or Fantasy? Neuron 70, 661–673 (2011).

14. Hill, R. A., Patel, K. D., Medved, J., Reiss, A. M. & Nishiyama, A. NG2 cells in white matter but not gray matter proliferate in response to PDGF. Journal of Neuroscience 33, 14558–14566 (2013).

15. Viganò, F., Möbius, W., Götz, M. & Dimou, L. Transplantation reveals regionaldifferences in oligodendrocytedifferentiation in the adult brain. Nat Neurosci 16, 1370–1372 (2013).

16. Falcão, A. M. et al. Disease-specific oligodendrocyte lineage cells arise in multiple sclerosis. Nat Med 24, 1837–1844 (2018).

17. Jäkel, S. et al. Altered human oligodendrocyte heterogeneity in multiple sclerosis. Nature 566, 543–547 (2019).

18. Spitzer, S. O. et al. Oligodendrocyte Progenitor Cells Become Regionally Diverse and Heterogeneous with Age. Neuron 101, 459–471 (2019).

19. Bergles, D. E., Roberts, J., Somogyl, P. & Jahr, C. Glutamatergic synapses on oligodendrocyte precursor cells in the hippocampus. Nature 405, 187–191 (2000).

20. De Biase, L. M., Nishiyama, A. & Bergles, D. E. Excitability and synaptic communication within the oligodendrocyte lineage. Journal of Neuroscience 30, 3600–3611 (2010).

21. Kukley, M., Capetillo-Zarate, E. & Dietrich, D. Vesicular glutamate release from axons in white matter. Nat Neurosci 10, 311–320 (2007).

22. Orduz, D. et al. Interneurons and oligodendrocyte progenitors form a structured synaptic network in the developing neocortex. Elife 4, e06953 (2015).

23. Káradóttir, R., Cavelier, P., Bergersen, L. H. & Attwell, D. NMDA receptors are expressed in oligodendrocytes and activated in ischaemia. Nature 438, 1162–1166 (2005).

24. Chittajallu, R., Aguirre, A. & Gallo, V. NG2-positive cells in the mouse white and grey matter display distinct physiological properties. The Journal of Physiology 561, 109–122 (2004).

25. Barres, B. A. & Raff, M. C. Proliferation of oligodendrocyte precursor cells depends on electrical activity in axons. Nature 361, 258–260 (1993).

26. Makinodan, M., Rosen, K. M., Ito, S. & Corfas, G. A Critical Period for Social Experience-Dependent Oligodendrocyte Maturation and Myelination. Science 337, 1357–1360 (2012).

27. Xiao, L. et al. Rapid production of new oligodendrocytes is required in the earliest stages of motor-skill learning. Nat Neurosci 19, 1210–1217 (2016).

28. Schebesta, M. & Serluca, F. C. olig1 Expression identifies developing oligodendrocytes in zebrafish and requires hedgehog and notch signaling. Dev. Dyn. 238, 887–898 (2009).

29. Auer, F., Vagionitis, S. & Czopka, T. Evidence for Myelin Sheath Remodeling in the CNS Revealed by In Vivo Imaging. Current Biology 28, 549–559.e3 (2018).

30. Picelli, S. et al. Smart-seq2 for sensitive full-length transcriptome profiling in single cells. Nat Meth 10, 1096–1098 (2013).

31. Emery, B. et al. Myelin gene regulatory factor is a critical transcriptional regulator required for CNS myelination. Cell 138, 172–185 (2009).

32. Chen, Y. et al. The oligodendrocyte-specific G protein-coupled receptor GPR17 is a cell-intrinsic timer of myelination. Nat Neurosci 12, 1398–1406 (2009).

33. Kirischuk, S., Scherer, J., Möller, T., Verkhratsky, A. & Kettenmann, H. Subcellular heterogeneity of voltage-gated Ca2+ channels in cells of the oligodendrocyte lineage. Glia 13, 1–12 (1995).

34. Pende, M., Holtzclaw, L. A., Curtis, J. L., Russell, J. T. & Gallo, V. Glutamate regulates intracellular calcium and gene expression in oligodendrocyte progenitors through the activation of DL-alpha-amino-3-hydroxy-5-methyl-4-isoxazolepropionic acid receptors. Proceedings of the National Academy of Sciences of the United States of America 91, 3215–3219 (1994).

35. Krasnow, A. M., Ford, M. C., Valdivia, L. E., Wilson, S. W. & Attwell, D. Regulation of developing myelin sheath elongation by oligodendrocyte calcium transients in vivo. Nat Neurosci 93, 1–28 (2017).

36. Leussink, V. I., Montalban, X. & Hartung, H.-P. Restoring Axonal Function with 4-Aminopyridine: Clinical Efficacy in Multiple Sclerosis and Beyond. CNS Drugs 32, 637–651 (2018).

37. Yu, X. et al. Reducing Astrocyte Calcium Signaling In Vivo Alters Striatal Microcircuits and Causes Repetitive Behavior. Neuron 99, 1–28 (2018).

38. Dimou, L. & Simons, M. Diversity of oligodendrocytes and their progenitors. Current Opinion in Neurobiology 47, 73–79 (2017).

39. Simons, M. & Lyons, D. A. Axonal selection and myelin sheath generation in the central nervous system. Current opinion in cell biology 25, 512–519 (2013).

40. Zhu, X. et al. Age-dependent fate and lineage restriction of single NG2 cells. Development 138, 745–753 (2011).

41. Hill, R. A., Patel, K. D., Goncalves, C. M., Grutzendler, J. & Nishiyama, A. Modulation of oligodendrocyte generation during a critical temporal window after NG2 cell division. Nat Neurosci 17, 1518–1527 (2014).

42. Bergles, D. E., Jabs, R. & Steinhäuser, C. Neuron-glia synapses in the brain. Brain research reviews 63, 130–137 (2010).

43. Kukley, M., Nishiyama, A. & Dietrich, D. The fate of synaptic input to NG2 glial cells: neurons specifically downregulate transmitter release onto differentiating oligodendroglial cells. Journal of Neuroscience 30, 8320–8331 (2010).

44. Wake, H. et al. Nonsynaptic junctions on myelinating glia promote preferential myelination of electrically active axons. Nat Commun 6, 7844 (2015).

45. Karttunen, M. J., Czopka, T., Goedhart, M., Early, J. J. & Lyons, D. A. Regeneration of myelin sheaths of normal length and thickness in the zebrafish CNS correlates with growth of axons in caliber. PLoS ONE 12, e0178058 (2017).

46. Almeida, R. G., Czopka, T., ffrench-Constant, C. & Lyons, D. A. Individual axons regulate the myelinating potential of single oligodendrocytes in vivo. Development 138, 4443–4450 (2011).

47. Freeman, J. et al. Mapping brain activity at scale with cluster computing. Nat Meth 11, 941–950 (2014).

48. Chen, T.-W. et al. Ultrasensitive fluorescent proteins for imaging neuronal activity. Nature 499, 295–300 (2013).

49. Bindels, D. S. et al. mScarlet: a bright monomeric red fluorescent protein for cellular imaging. Nat Meth 14, 53–56 (2017).

50. Almeida, R. G. & Lyons, D. A. Intersectional Gene Expression in Zebrafish Using the Split KalTA4 System. Zebrafish 12, 377–386 (2015).

51. Kwan, K. M. et al. The Tol2kit: a multisite gateway-based construction kit for Tol2 transposon transgenesis constructs. Dev. Dyn. 236, 3088–3099 (2007).

52. Walton, E. M., Cronan, M. R., Beerman, R. W. & Tobin, D. M. The Macrophage-Specific Promoter mfap4 Allows Live, Long-Term Analysis of Macrophage Behavior during Mycobacterial Infection in Zebrafish. PLoS ONE 10, e0138949 (2015).

53. Czopka, T., ffrench-Constant, C. & Lyons, D. A. Individual oligodendrocytes have only a few hours in which to generate new myelin sheaths in vivo. Developmental Cell 25, 599–609 (2013).

54. Martin, M. Cutadapt Removes Adapter Sequences From High-Throughput Sequencing Reads. EMBnet.journal 17, 3 (2011).

55. Dobin, A. et al. STAR: ultrafast universal RNA-seq aligner. Bioinformatics 29, 15–21 (2013).

56. Patro, R., Duggal, G., Love, M. I., Irizarry, R. A. & Kingsford, C. Salmon provides fast and bias-aware quantification of transcript expression. Nat Meth 14, 417–419 (2017).

57. Butler, A., Hoffman, P., Smibert, P., Papalexi, E. & Satija, R. Integrating single-cell transcriptomic data across different conditions, technologies, and species. Nature biotechnology 36, 411–420 (2018).

58. Bindea, G. et al. ClueGO: a Cytoscape plug-in to decipher functionally grouped gene ontology and pathway annotation networks. Bioinformatics 25, 1091–1093 (2009).

59. Fiji: an open-source platform for biological-image analysis. Nat Meth 9, 676–682 (2012).

60. Meijering, E. et al. Design and validation of a tool for neurite tracing and analysis in fluorescence microscopy images. Cytometry A 58, 167–176 (2004).

