## Supplementary material for "Functionally Distinct Subgroups of Oligodendrocyte Precursor Cells Integrate Neural Activity and Execute Myelin Formation": Hoche Marisca et al supplementary figures

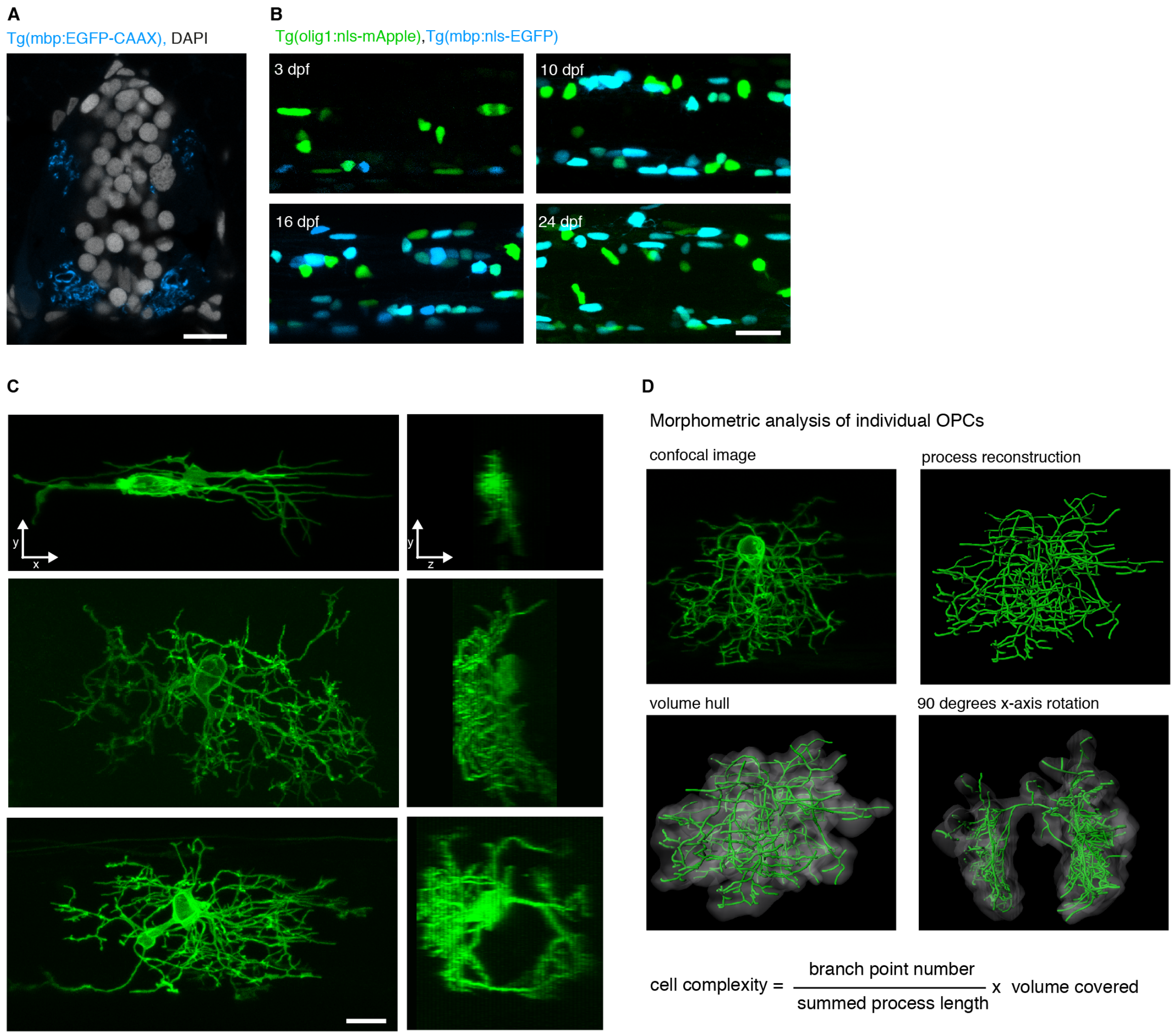

**Figure S1: Supplementary to Figure 1**

**(A)** Transversal section of a Tg(mbp:EGFP-CAAX) zebrafish at 7dpf showing the position of myelinated axons in the spinal cord. Scale bar: 10 $\mu$m

**(B)** Confocal images of Tg(olig1:nls-mApple), Tg(mbp:nls-EGFP) spinal cord at different ages (3 dpf, 10 dpf, 16 dpf, 24 dpf) to supplement images shown in Fig. 1C. Scale bar: 20 $\mu$m.

**(C)** Confocal images of individual OPCs labelled by injection of olig1:memYFP showing the range different morphologies. The soma can be localized in the spinal cord white (top) or grey matter (middle, bottom). The process network of an individual cell can be restricted to one the side of the spinal cord (top and middle cells), but can also reach to the opposite site of the spinal cord (bottom cell). Scale bar: 10 $\mu$m.

**(D)** Example images to show how morphometry of individual cells was conducted. For a three-dimensional confocal image (top left) all the processes for an individual cell were traced (top right). To get a measure for the volume a hull was constructed around the reconstructed filament network (bottom).

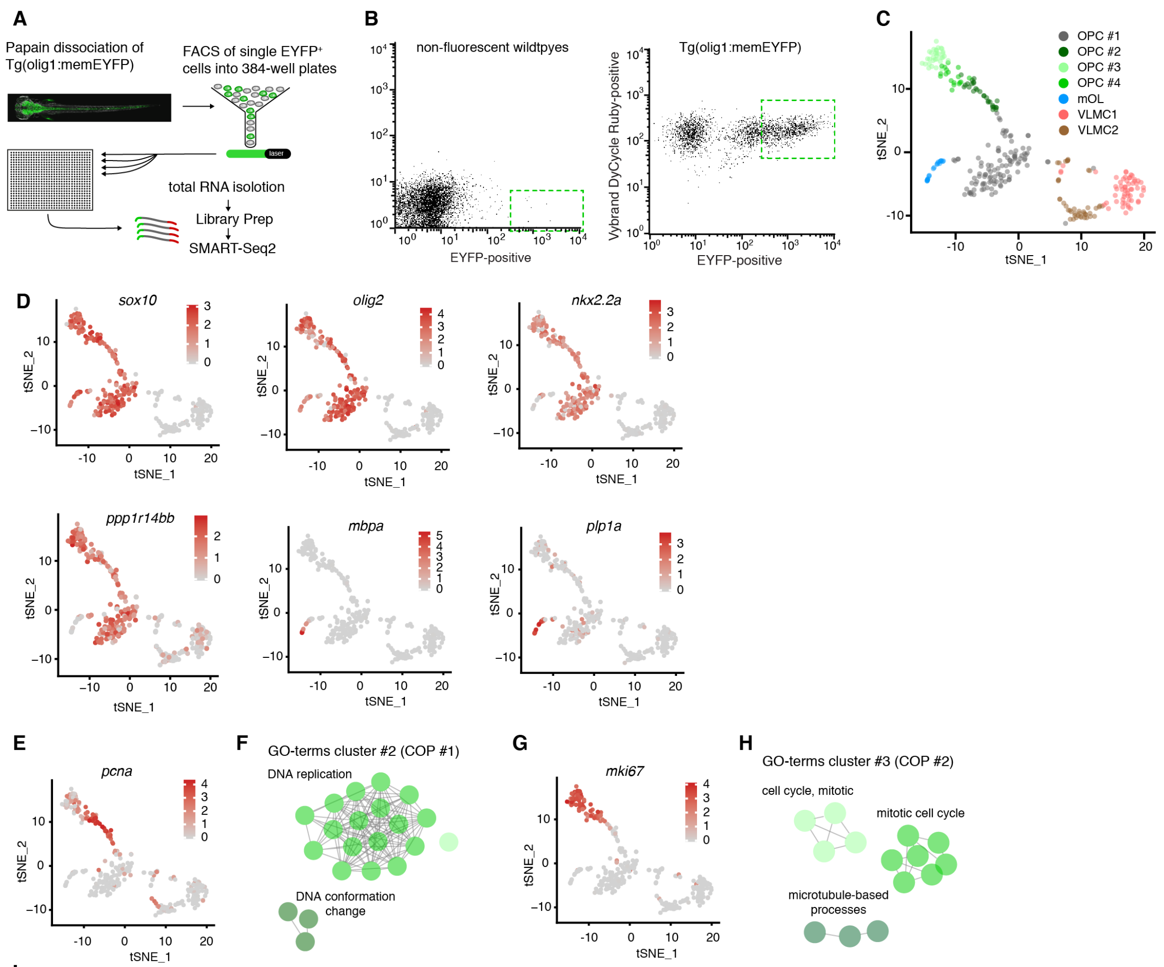

**I**

| OPC #1 | | |
|---|---|---|
| gene name | avg_logFC | p-value |
| glula | 1,66 | 3,92E-28 |
| aplnrb | 1,58 | 1,80E-28 |
| gria1a | 1,50 | 6,77E-30 |
| slc1a2b | 1,48 | 3,06E-25 |
| gria2b | 1,47 | 1,33E-30 |
| atcaya | 1,42 | 3,76E-29 |
| wscd1b | 1,41 | 3,57E-24 |
| ncalda | 1,36 | 2,08E-24 |
| mdkb | 1,36 | 9,95E-28 |
| gpm6aa | 1,36 | 2,39E-31 |
| scg2b | 1,35 | 1,29E-27 |
| slc6a1a | 1,33 | 1,34E-24 |
| nrxn1a | 1,32 | 2,07E-24 |
| pcdh2ab9 | 1,31 | 1,81E-25 |
| ckbb | 1,31 | 2,66E-28 |
| grin1b | 1,31 | 3,52E-24 |
| tmem178b | 1,30 | 5,49E-28 |
| camk2n1a | 1,30 | 1,62E-27 |
| olig1 | 1,29 | 4,06E-24 |
| nlgn3a | 1,29 | 3,46E-27 |
| cspg5a | 1,26 | 2,50E-27 |
| sema5a | 1,25 | 6,44E-28 |
| lrrn1 | 1,22 | 2,75E-22 |
| fabp7a | 1,22 | 3,72E-24 |
| ndrg3a | 1,21 | 3,51E-25 |
| rassf2a | 1,21 | 3,74E-26 |
| ttyh3b | 1,20 | 6,44E-25 |
| spock3 | 1,18 | 1,51E-26 |
| kiaa1549la | 1,18 | 9,75E-24 |
| pygmb | 1,17 | 1,86E-23 |
| cspg4 | 0,80 | 2,15E-13 |

| OPC #2 | | |
|---|---|---|
| gene name | avg_logFC | p-value |
| pcna | 2,72 | 1,38E-20 |
| rpa1 | 1,99 | 5,62E-22 |
| slc29a2 | 1,96 | 9,20E-17 |
| fen1 | 1,84 | 7,88E-26 |
| rrm2 | 1,83 | 6,03E-19 |
| dnmt1 | 1,78 | 7,24E-19 |
| zgc:110540 | 1,72 | 1,92E-25 |
| stmn1a | 1,63 | 9,53E-16 |
| mcm5 | 1,60 | 6,46E-09 |
| dnajc9 | 1,60 | 2,35E-18 |
| rrm1 | 1,58 | 2,97E-23 |
| rbbp4 | 1,53 | 3,34E-16 |
| slbp | 1,52 | 5,10E-31 |
| lig1 | 1,50 | 6,27E-26 |
| rrm2 | 1,47 | 9,29E-14 |
| rpa2 | 1,44 | 3,26E-22 |
| mcm2 | 1,44 | 4,20E-12 |
| chaf1a | 1,41 | 4,54E-22 |
| nasp | 1,41 | 2,64E-18 |
| atad2 | 1,40 | 1,41E-21 |
| mcm3 | 1,40 | 4,82E-18 |
| mibp | 1,33 | 2,78E-10 |
| rfc5 | 1,32 | 1,33E-17 |
| mcm4 | 1,30 | 2,67E-08 |
| sumo3b | 1,29 | 1,29E-14 |
| tubb2b | 1,28 | 2,35E-12 |
| asf1ba | 1,21 | 1,97E-12 |
| dck | 1,20 | 3,25E-18 |
| atad5a | 1,20 | 1,05E-20 |
| cdca7a | 1,20 | 5,14E-11 |

| OPC #3 | | |
|---|---|---|
| gene name | avg_logFC | p-value |
| kpna2 | 2,86 | 1,92E-23 |
| cenpf | 2,62 | 1,50E-27 |
| aspm | 2,55 | 3,40E-28 |
| tuba8l | 2,48 | 2,53E-16 |
| kif11 | 2,45 | 1,41E-29 |
| plk1 | 2,36 | 5,53E-27 |
| ccnb1 | 2,23 | 7,20E-31 |
| tpx2 | 2,21 | 2,10E-31 |
| top2a | 2,17 | 1,68E-25 |
| cdk1 | 2,16 | 1,42E-26 |
| nusap1 | 2,15 | 1,26E-28 |
| ube2c | 2,10 | 4,63E-29 |
| mki67 | 2,10 | 2,11E-26 |
| tacc3 | 2,08 | 1,44E-34 |
| cdc20 | 2,06 | 8,57E-36 |
| cdc14b | 2,03 | 6,80E-23 |
| prc1b | 1,94 | 4,04E-34 |
| kifc1 | 1,92 | 2,26E-29 |
| prc1a | 1,92 | 1,75E-27 |
| g2e3 | 1,91 | 1,28E-30 |
| ccna2 | 1,86 | 9,00E-25 |
| cenpe | 1,77 | 3,17E-29 |
| cdca8 | 1,75 | 2,37E-30 |
| ccnf | 1,74 | 2,36E-31 |
| anln | 1,73 | 5,74E-31 |
| mastl | 1,73 | 2,08E-28 |
| mad2l1 | 1,71 | 2,13E-31 |
| ccnb3 | 1,71 | 9,96E-33 |
| rtkn2a | 1,70 | 2,49E-31 |
| kif2c | 1,70 | 6,93E-27 |

| OPC #4 | | |
|---|---|---|
| gene name | avg_logFC | p-value |
| gpr17 | 1,94 | 8,34E-25 |
| prdm8 | 1,85 | 4,93E-13 |
| draxin | 1,82 | 4,46E-12 |
| hmgn2 | 1,81 | 1,55E-11 |
| hmgb2a | 1,70 | 6,68E-11 |
| lmnb1 | 1,67 | 7,53E-12 |
| cdk1 | 1,60 | 7,18E-13 |
| mki67 | 1,56 | 4,63E-13 |
| top2a | 1,52 | 5,23E-11 |
| plk1 | 1,50 | 5,03E-12 |
| aspm | 1,50 | 6,28E-13 |
| hmga1a | 1,50 | 3,68E-10 |
| lbr | 1,47 | 3,99E-12 |
| nusap1 | 1,45 | 3,37E-13 |
| ccnb1 | 1,45 | 3,09E-14 |
| sox9b | 1,44 | 3,71E-10 |
| tuba8l4 | 1,44 | 9,82E-11 |
| cenpe | 1,38 | 4,52E-15 |
| hmgb2b | 1,36 | 9,28E-11 |
| kpna2 | 1,35 | 4,94E-09 |
| zeb2a | 1,34 | 9,61E-11 |
| tnr | 1,34 | 9,69E-09 |
| marcksb | 1,32 | 8,60E-11 |
| cenpf | 1,31 | 1,47E-11 |
| sox8b | 1,30 | 8,16E-18 |
| kif11 | 1,30 | 1,19E-12 |
| mycb | 1,29 | 1,76E-13 |
| tuba8l | 1,27 | 7,19E-08 |
| wu:fb44b02 | 1,27 | 7,67E-16 |
| tpx2 | 1,27 | 2,35E-13 |
| myrf | 0,97 | 1,96E-10 |

### Figure S2

**Figure S2: Supplementary to Figure 2**

**A)** Schematic overview of cell isolation, sorting and sequencing.

**B)** Flow cytometry plots of olig1:memEYFP-sorted cells and wildtype control animals.

**C)** TSNE plots showing clusters identified from all sequenced olig1:memEYFP cells.

**D)** TSNE plots with gene expression levels of different oligodendrocyte markers from all sequenced olig1:memEYFP cells.

**E)** TSNE plots with *pcna* gene expression levels from all sequenced olig1:memEYFP cells.

**F)** GO terms of top 30 differentially expressed genes in cluster OPC #2.

**G)** TSNE plots with *mki67* gene expression levels from all sequenced olig1:memEYFP cells.
**H)** GO terms of top 30 differentially expressed genes in cluster OPC #3.
**I)** Lists of top 30 differentially expressed genes in OPC clusters #1 to #4.

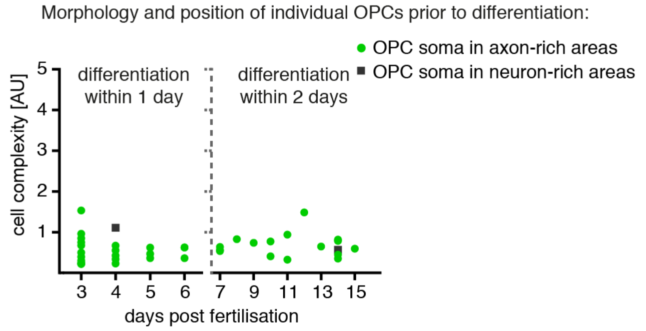

**Figure S3: Supplementary to Figure 3**
**A)** Quantification of position and morphology of individual OPCs from imaging timelines between 3 and 15 dpf (n=43 cells). Measured is the last timepoint prior to differentiation, as assessed by myelin sheath formation (imaging intervals of 1d between 3 and 7 dpf, and 2d between 7 and 15dpf).

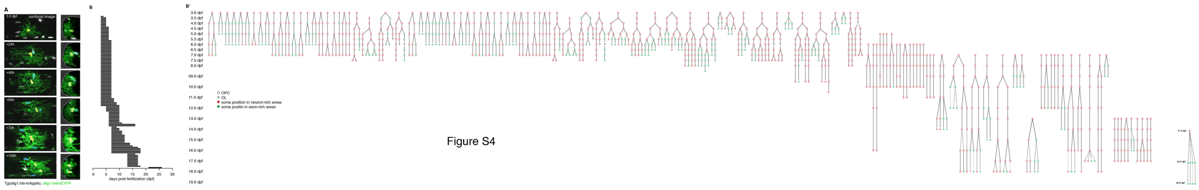

**Figure S4: Supplementary to Figure 4**
A) Projections (left) and 90˚ y-axis rotations (right) of timelines of confocal images of individual olig1:memEYFP labelled cell with its soma in neuron-rich areas in Tg(olig1:mls-mApple) analysed in Figure 4A. Scale bar: 10 $\mu$m.
**B)** Overview of time windows during which clonal OPC fates were analysed. Analysis started at different time points post fertilization. B') For each tree, individually labelled olig1:memEYFP cells positioned with their soma in neuron-rich areas were selected as starting point and followed for at least four days or until the cell or respective daughter cell differentiated.

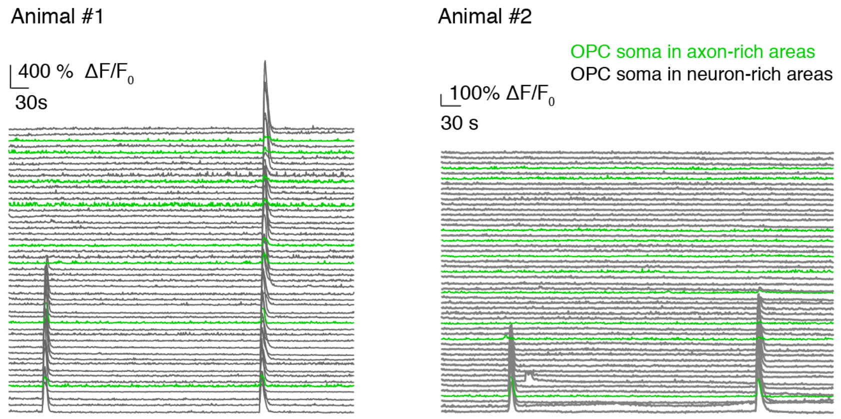

**Figure S5: Supplementary to Figure 5**
ΔF/F0 GCaMP transients of individual cells in Tg(olig1:GCaMP6m). Green traces depict cells in axon-rich areas, grey traces depict cells in neuron-rich areas.

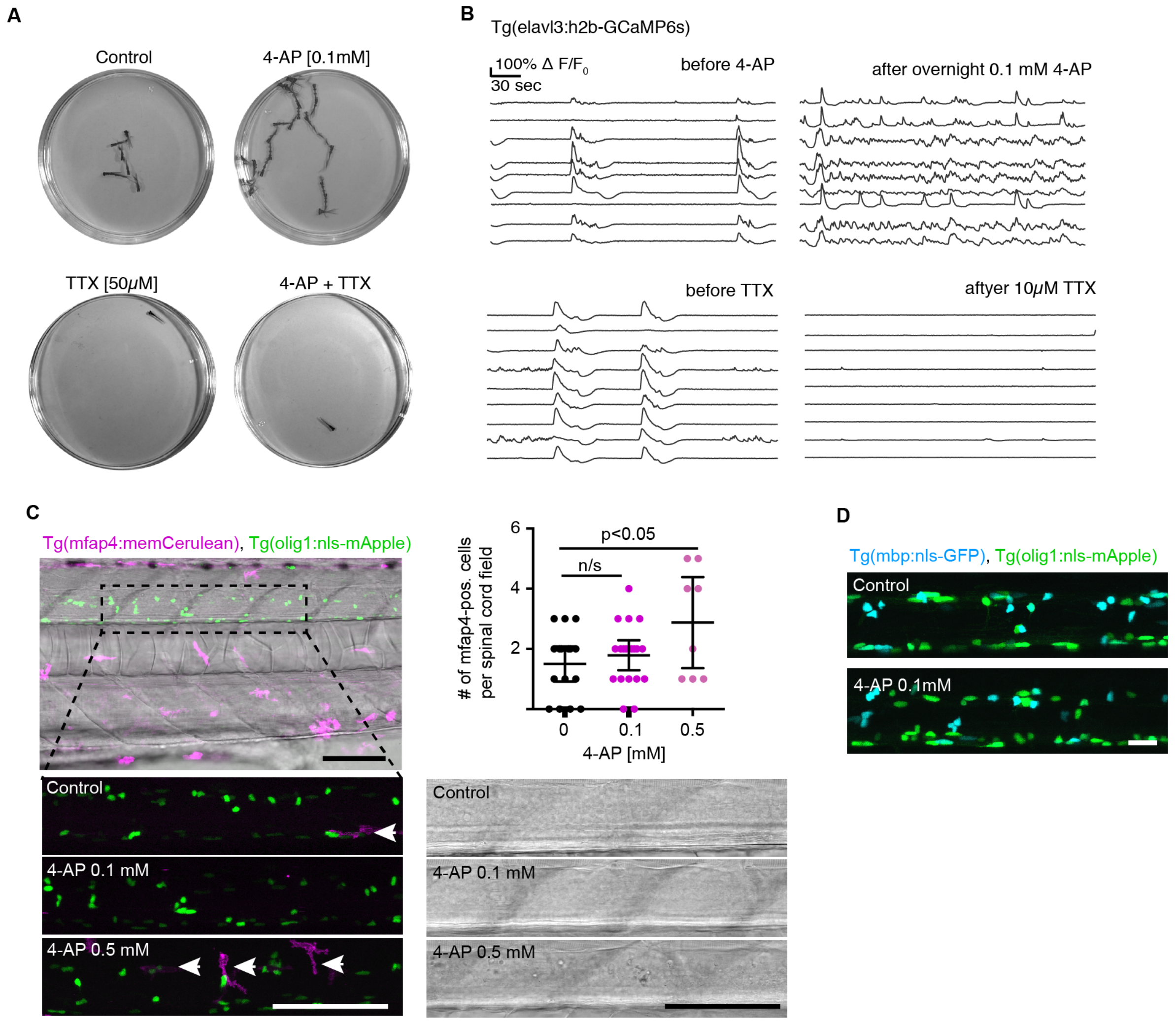

**Figure S6. Supplementary to Figure 6**

**A)** Minimum intensity projections of a two-minute time-lapse of fish freely swimming in a 3cm petri dish in different treatment conditions.

**B)** Traces of GCaMP transients Tg(elavl3:h2b-GCaMP6s) zebrafish at 4dpf and after overnight incubation in 0.1mM 4-AP, and before / after 10 μM TTX.

**C)** Confocal images of Tg(mfap4:memCerulean), Tg(olig1:nls-mApple) zebrafish at 4dpf after treatment with 0.1mM or 0.5mM 4-AP or Danieau's solution as control. Transmitted light images to show spinal cord morphology and tissue integrity following drug treatment. Scale bars: 100 μm. The graph shows that number of macrophages which accumulate in 400 μM length of spinal cord of Tg(mfap4:memCerulean) zebrafish after 1 day of control, 0.1 mM or 0.5 mM 4-AP treatment. Data are expressed at mean ± 95% CI, n=16/19/8 animals (control/0.1 mM 4-AP / 0.5 mM 4-AP), control *vs.* 0.5 mM 4-AP, p=0.03 (one-way ANOVA).

**D)** Confocal images Tg(mbp:nls-EGFP), Tg(olig1:nls-mApple) zebrafish in control and following 2 days of 0.1mM 4-AP treatment. Scale bar: 20 μm.

**Movie S1: Pan and zoom of Tg(olig1:memEYFP).**

**Movie S2: Segmentation of individual OPCs.**

**Movie S3: 60 minutes time-lapse of individual OPC with its soma in axon-rich areas.**

**Movie S4: 60 minutes time-projection of individual OPC with its soma in axon-rich areas.**

**Movie S5: 60 minutes time-lapse of individual OPC with its soma in neuron-rich areas.**

**Movie S6: 60 minutes time-projection of individual OPC with its soma in neuron-rich areas.**

**Movie S7: 24 hours time-lapse of individual OPC with its soma in neuron-rich areas.**

**Movie S8: 13 hours time-lapse of individual OPC with its soma in axon-rich areas.**

**Movie S9: Time-lapse of two olig1-GCaMP6m-CAAX labelled OPCs showing transients in process subdomains.**

**Movie S10: Time-lapse of two olig1-GCaMP6m-CAAX labelled OPCs showing transients throughout cell.**

**Movie S11: Time-lapse of a Tg(olig1:GCaMP6m) at the level of the spinal cord (dorsal view).**

**Table S1: Primers used for molecular cloning**

| Primer name | Sequence |
|---|---|
| attB1_GCaMP6m_F | GGGGACAAGTTTGTACAAAAAAGCAGGCTGCCACCATGGGTTCTCATC |
| attB2R_GCaMP6m_R | GGGGACCACTTTGTACAAGAAAGCTGGGTCTCACTTCGCTGTCATCATTTGTA |
| CAAX-GCaMP6m_R | TCAGGAGAGCACACACTTGCAGCTCATGCAGCCGGGGCCACTCTCATCAGGAGGGTTCAGCTTTCACTTCGCTGTCATCATTTGTAC |
| attB2R_CAAX_R | GGGGACCACTTTGTACAAGAAAGCTGGGTCTCAGGAGAGCACACACTTGC |
| attB1_mCherry_F | GGGGACAAGTTTGTACAAAAAAGCAGGCTGCCACCATGGTGAGCAAGGGCGAG |
| attB2R_CalEx-R | GGGGACCACTTTGTACAAGAAAGCTGGGTCCTAAAGCGACGTCTCCAG |
| attB4_mfap4 | GGGGACAACTTTGTATAGAAAAGTTGGCGTTTCTTGGTACAGCTGG |
| attB1R_mfap4 | GGGGACTGCTTTTTTGTACAAACTTGCTTCTCACTCTCTCCTCAAC |

Colour code: att recombination site, Kozak sequence, coding sequence

**Table S2: Antibodies and dyes**

| Name | Organism / type | Company |
|---|---|---|
| anti-DSRed | polyclonal rabbit IgG | Takara/Clontech |
| anti-mCherry | polyclonal chicken IgY | Novus Biologicals |
| anti-GFP | polyclonal chicken IgY | Abcam |
| 3A10 | monoclonal mouse IgG1 | Developmental Studies Hybridoma Bank |
| anti Sox10 | polyclonal rabbit IgG | BIOZOL |
| anti-chicken Alexa Fluor 488 | polyclonal goat IgG | Invitrogen |
| anti-chicken Alexa Fluor 555 | polyclonal goat IgG | Invitrogen |
| anti-rabbit AlexaFluor 555 | polyclonal goat IgG | Invitrogen |
| anti-rabbit Alexa Fluor 633 | polyclonal goat IgG | Invitrogen |
| anti-mouse Alexa Fluor 633 | polyclonal goat IgG | Invitrogen |
| BODIPY 630/650-X | n/a | Thermo Fisher |
| Opal 650 | n/a | Perkin Elmer |
